# High-quality complete genome resource of tomato rhizosphere strain *Pseudomonas donghuensis* P482, a representative of a species with biocontrol activity against plant pathogens

**DOI:** 10.1101/2021.06.09.447503

**Authors:** Dorota M. Krzyżanowska, Adam Iwanicki, Robert Czajkowski, Sylwia Jafra

## Abstract

Strain P482 was isolated from a tomato rhizosphere and classified as *Pseudomonas donghuensis*. The *P. donghuensis* species was first established in 2015 and currently comprises only four strains: P482, HYS^T^, SVBP6, and 22G5. *P. donghuensis* strains antagonize plant pathogens, including bacteria, fungi and oomycetes, and therefore are of high interest regarding their biological control potential to combat plant diseases. The antimicrobial activity of *P. donghuensis* P482 is based on the production of iron scavenging compound 7-hydroxytropolone, antifungal volatile organic compounds, and yet unidentified secondary metabolite(s). Here, we report a complete genome resource for *P. donghuensis* strain P482. The genome consists of a single chromosome (5 656 185 bp) with 5258 ORFs (5158 protein-coding genes, 74 tRNAs, 22 rRNAs, 3 ncRNAs and 1 tmRNA) and no plasmid. We believe that the information of the first high-quality, complete genome of *P. donghuensis* will provide resources for analyses targeting the biological control potential of this species and understanding the traits essential for plant-microbe interaction.

## Genome Announcement

*Pseudomonas* spp. constitute a group of Gammaproteobacteria known for their ability to adapt to various environments, metabolize multiple carbon sources, and produce a remarkable array of secondary metabolites (Gross and Loper, 2009). Although the genus contains pathogens such as *P. aeruginosa, P. entomophila* or *P. syringe*, most *Pseudomonas* spp. are harmless dwellers or beneficial organisms(Peix et al., 2018). Many representatives of the genus promote plant growth and/or protect their hosts from the harmful activity of plant pathogens, placing them within the group of the Plant Growth Promoting Rhizobacteria (PGPR) (Lugtenberg and Kamilova, 2009). Significantly, the mechanism of PGPR activity of pseudomonads can differ depending on a species or even a particular strain – a feature related to the high versatility of genome size and content among *Pseudomonas* spp. (Silby et al., 2011; Loper et al., 2012). Due to their PGPR properties, the plant-beneficial *Pseudomonas* spp. are of high interest in application as biological plant protection agents in integrated farming (Mercado-Blanco, 2015).

In our previous studies, we isolated the P482 strain from the rhizosphere of a tomato plant grown in a garden in Gdynia, Poland (Krzyzanowska et al., 2012). The P482 inhibits the growth of bacterial plant pathogens of the soft rot *Pectobacteriaceae* (SRP) (former pectinolytic *Erwinia* spp.) group, as well as *Pseudomonas syringae*, and it attenuates disease symptoms caused by these pathogens on plant tissues (Golanowska et al., 2012; Krzyzanowska et al., 2012). Moreover, volatile organic compounds (VOCs) produced by P482 can inhibit the growth of fungal pathogens *Rhizoctonia solani*, *Fusarium culmorum, Verticillium dahliae* and oomycete *Phytium ultimum* (Ossowicki et al., 2017). The phylogenetic studies revealed that the strain belongs to *P. donghuensis* – a species represented exclusively at that time by one strain: HYS^T^ (Krzyzanowska et al., 2016). Currently, the *P. donghuensis* species comprises four strains: P482, HYS^T^, SVBP6, and 22G5 (Gao et al., 2015; Krzyzanowska et al., 2016; Agaras et al., 2018; Tao et al., 2020), all of which were shown to have potential to antagonize plant pathogens (Krzyzanowska et al., 2012; Yu et al., 2014; Agaras et al., 2015; Tao et al., 2020). The antimicrobial properties of *P. donghuensis* strains are attributed mainly, but not exclusively, to the production of antimicrobial and iron scavenging compound 7-hydroxytropolone (7-HT). 7-HT is not a typical metabolite of *Pseudomonas* spp. The *nfs* gene cluster necessary for its production has, so far, been detected solely in *P. donghuensis* (Yu et al., 2014; Krzyzanowska et al., 2016; Muzio et al., 2020; Tao et al., 2020). Prior detection of 7-HT in *P. donghuensis*, the production of this metabolite was reported in *Pseudomonas lindbergii* ATCC31099 (Korth et al., 1981) and *Streptomyces neyagawaensis* (Kirst et al., 1982). In the latter, it was studied for its ability to decrease microbial resistance to aminoglycoside antibiotics by the inhibition of aminoglycoside adenyltransferase (Allen et al., 1982). Nonetheless, P482 is also known to produce antifungal VOCs, and, as we have recently unveiled, its genome harbors gene clusters involved in the synthesis of yet unidentified secondary metabolites (Matuszewska et al., 2021).

Until now, none of the *P. donghuensis* genomes has been delivered in the complete (closed) form. Herein, for the first time, we provide a complete, high-quality genome sequence of *P. donghuensis* strain P482.

Draft genome of strain P482 was released in 2014 (Krzyzanowska et al., 2014). At that time, the quality of data and the available software did not allow the assembly of reads into a single molecule, resulting in 69 contigs. Sequencing of genomes of the three other representatives of *P. donghuensis* also did not yield a closed chromosome (Table 1). In this study, to refine the genomic sequence of *P. donghuensis* P482, the strain was cultured for 48 hours on LB agar medium. Isolation of DNA and sequencing was performed at Oligo.pl (Institute of Biochemistry and Biophysics, Polish Academy of Sciences, Warsaw, Poland). Genome sequencing was performed using two sequencing platforms: Illumina MiSeq (69 × coverage) and Oxford Nanopore GridION (109 × coverage). Hybrid assembly of quality filtered reads was performed using Unicycler 0.4.8 (Wick et al., 2017). The remaining ambiguities and sequence errors were verified by PCR and Sanger sequencing. The resulting high-quality complete genome of strain P482 contains one circular chromosome of 5 656 185 bp with an average G+C content of 62.36%. No plasmid was detected. The sequence was annotated using the NCBI Prokaryotic Genome Annotation Pipeline (PGAP), version March 2021 (Li et al., 2021). The genome was found to contain 5 258 ORFs, among which there are 5 158 CDSs (CoDing Sequences), 74 tRNAs, 22 rRNAs, 3 ncRNAs and 1 tmRNA.

**Table 1.** Strains of *P. donghuensis* and the level of assembly of available genomic data.

| Strain | Strain | Assembly | GenBank | Total | G+C | ANI | Reference |
| --- | --- | --- | --- | --- | --- | --- | --- |
|  | isolation | level | accession | sequence | % | %<sup>A,B</sup> | (genome) |
|  | source | (No. of |  | length |  |  |  |
|  |  | contigs) |  |  |  |  |  |
| P482 | Rhizosphere of tomato, Poland | Complete (1) | CP071706 | 5 656 185 | 62.36 | 99.47<sup>B</sup> | This study |
|  |  | WGS (69) | JHTS000000000.1 | 5 623 997 | 62.38 | 99.47 | (Krzyzanowska et al., 2014) |
| HYS<sup>T</sup> | Lake water, China | WGS (231) | AJJP000000000.1 | 5 646 028 | 62.42 | 100 | (Gao et al., 2012) |
| SVBP6 | Bulk soil of an agricultural plot, Argentina | WGS (43) | NWCB000000000.1 | 5 701 342 | 62.37 | 99.45 | (Agaras et al., 2018) |
| 22G5 | Rhizosphere of weeds, China | WGS (3185) | RWIB000000000.1 | 6 546 541 | 60.74 | 99.41 | (Tao et al., 2020) |

<sup>A</sup> ANI calculated against type strain HYS<sup>T</sup> using ChunLab's online Average Nucleotide Identity (ANI) calculator (https://www.ezbiocloud.net/tools/ani) (Yoon et al., 2017)
<sup>B</sup> ANI calculated for the complete chromosome of P482 and the draft version from 2014 equaled 99.98%
<sup>T</sup> type strain for the species

New annotation performed for the complete genome provides us with the advantage of up-to-date annotation of protein functions. However, studies referring to the four-digit locus tags in the draft genome of P482 have already been published (Krzyzanowska et al., 2016; Ossowicki et al., 2017; Matuszewska et al., 2021 - *in press*). To avoid confusion regarding the designations of genes, we performed cross-mapping of CDSs between the draft and the newly-annotated complete genome reported in this study. The mapping was performed at the protein level using BLASTP 2.11.0+ (Altschul et al., 1997) (Supplementary Materials File 1). The threshold for successful mapping was set to E-value ≤0.001 and identity ≥96%. As a result, locus tags for 4942 protein coding genes (95,8%) were transferred from the draft to the complete genome. Proteins encoded by 4433 mapped CDSs showed the identity of 100% between versions, 487 showed identity of 99% and 22 showed identity between 96-98%. All remaining genes, including non-protein coding genes, were designated with new five-digit locus tags.

Comparison of the refined sequence of *P. donghuensis* P482 with the draft revealed that the complete chromosome contains 32 188 base pairs more than the draft. Figure 1. depicts the order and orientation of genome content in the chromosome in relation to the succeeded WGS assembly. The analysis was performed using progressiveMauve (Darling et al., 2010). The same program revealed 519 single nucleotide polymorphisms (single base changes) between the complete genome and the draft (Supplementary Materials File 2). The cross-mapping of loci revealed 188 protein coding sequences from the draft that could not be mapped to the complete chromosome of P482 (Supplementary Materials File 3). Most of the unmapped genes encoded either hypothetical of putative proteins (92% in total) and only 4 (2.1%) of the remaining entries could be assigned function by KEGG using BlastKOALA (Kanehisa et al., 2016). On the opposite, 118 protein coding sequences present in the new annotation of the closed genome could not be mapped to the draft (Supplementary Materials File 4). Out of this set 11 entries (13%) were assigned a KEGG KO identifier. Among the latter were such genes as *recC, recD* and *pqqA*. The differences in detected ORFs between the complete genome and the draft result from the differences in the sequence, the fact that some ORFs were not identified in the draft due to its fragmentation into 69 contigs, as well as from applying another, and more up-to-date engine to annotate the genome of P482.

**Figure 1.**
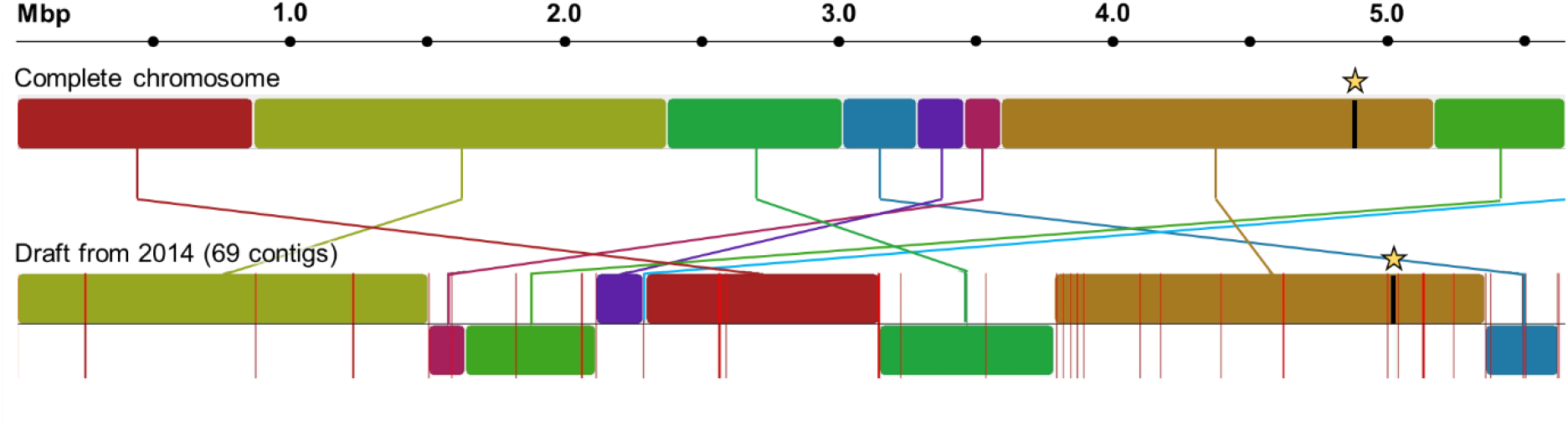
The order and orientation of genome content in the complete chromosome of *P. donghuensis* P482 (this study) in relation to the superseded draft genome from 2014. Analysis was performed using progressiveMauve. Each color block represents a region of the complete genome of P482 that aligned to the WGS version. Sequences within the blocks are presumed to be free from internal genomic rearrangements. Placement of the blocks below the centerline indicates that a particular region aligns in the reverse complement orientation. Red vertical lines in the draft version represent contig borders. Stars indicate the genomic location of the homologue of the *nfs* gene cluster encoding the production of 7-hydroxytropolone – an important antimicrobial compound produced by the strains of *P. donghuensis*.

In our previous study, the draft genome sequence of P482 was analyzed for the presence of genes encoding the production of secondary metabolites using antiSMASH 2.0 (Blin et al., 2013). The program revealed the presence of 23 gene clusters, including 5 with any suggestion of their function (two non-ribosomal peptide synthases (“NRPS”), one “Bacteriocin”, and two “Other”) and 18 putative clusters, one of which was found to contain the *nfs* genes required to produce 7-HT (Krzyzanowska et al., 2016). Here, we analyzed the complete genome of P482 with antisMASH 6.0 beta (Blin et al., 2021) in similar setting allowing the most extended search. The updated version of antiSMASH found genomic regions resembling those involved in the synthesis of metabolites such as pseudopyronine (62%), aryl polyen Vf (40%), fragin (37%), lipopolysaccharides (27%, 5%), O-antigens (19% and 14%), pyoverdine (17%, 11%, 6%), lankacidin C (13%) and chejuenolide (7%). Interestingly, however, the putative clusters predicted by antiSMASH 6.0 did not contain region now known to encode the synthesis of 7-HT.

The complete genome of P482 was investigated for the presence of prophage sequences using Prophage Hunter (https://pro-hunter.genomics.cn) (Song et al., 2019). The analysis resulted in discovering a single putatively intact (active) prophage genome of 57 737 bp, located between positions 2812547 and 2870283. This prophage genome was shown to contain genes vital for temperate bacteriophages. These included: integrase, regulators governing switch between lytic and lysogenic life cycle, genes coding for holin and lysin and structural proteins. According to PHASTER (Zhou et al., 2011), this prophage is phylogenetically the most similar to *Pseudomonas* phage YMC/01/01/P52_PAE_BP (Jeon et al., 2012). The presence of structural genes in the prophage genome was verified by the VirFam (http://biodev.cea.fr/virfam/) (Lopes et al., 2014). It allowed classification of the prophage into order *Caudovirales* and family *Siphoviridae*.

Genome resource for *P. donghuensis* P482 described in this study was deposited in GenBank under accession number CP071706.

## Authors’ statement

The authors declare no conflict of interest.

## Acknowledgements

The work was financially supported by the Polish National Science Centre (Narodowe Centrum Nauki, Polska) research grant OPUS13 no. 2017/25/B/NZ9/00513 and Polish Ministry of Science and Higher Education (Ministerstwo Nauki i Szkolnictwa Wyższego, Polska) funds DS 531-M105-D786-20 to SJ and by Polish Ministry of Education and Science (Ministerstwo Szkolnictwa I Nauki, Polska) funds DS 531-N104-D800-21 to RC.

## Notes

### Competing Interest Statement

The authors have declared no competing interest.

