## Supplementary Materials 3 & 4 for "High-quality complete genome resource of tomato rhizosphere strain *Pseudomonas donghuensis* P482, a representative of a species with biocontrol activity against plant pathogens"

**Supplementary File 3**
**188 Frames from draft not mapped to complete genome of P482.**
>KDN96928.1 hypothetical protein BV82_5230C, partial [Pseudomonas donghuensis]
EVKLAAKFCVASVASTVSSTVSVGSVPVVLMLGPSSSKIIFEPISAVVVEAWSSLKPPISKSAWLLPLAS
TAAPFSMSTRLWSAVVVYSIL

>KDN96929.1 hypothetical protein BV82_5231 [Pseudomonas donghuensis]
MVCRCGVFNPVAGFNAGDGRNPGSKADDNGRIIRRSVRVLDHQPGADVDDANIADCNRRITHIAVDEIRA
RFSSGAECADFYHRALAALQLHDVERGRPRIEDIGEVDVGFNICFVLIQVGNGHILPGTGDERKADALLF
NNHLVVSAEGGRATCLAGIDRIARRYIGAARLVGGVLAGGDNGRVSDPQDPTVVDHGRVRRREHQEQVLP
GRRIGQAEYQVLAGADEHLVQQVEGIVRVTGVWMLERLVFHGPDLAAGDRRSAFLNVGREHDRTGWPPGS
AQQDVVVGIGIKKDAGAIEGIESAVQAIERCAQAVGEADIGNAGSGCRGDGERIAAGVTHRPRIATQGCT
CCGTGPQRCQAGSEGRGQRRDQNVIVVSVSRRACGGRAVLQSGGFVQRNVHPRRRDTQGAGHGDGSAGGG
GVCRVGGGDRDGKQAGISVIGKVARRRCGFFADCHSVLPSKPSHRGWSVDLYC

>KDN96933.1 hypothetical protein BV82_5230B, partial [Pseudomonas donghuensis]
DLAPGHRQLHRIDQRRALLLNEEGYLPTGLRRQGVDLDEVRECAILVVDPVVQVLGRRVVSPHGVVDEHL
HGCQQALRVDDETFIGVADGCQVVPLEEVIGDADAGCADVHGRRVYVRWQGVGLQRSFSGVGFSEPDAGQ
DVRRRRKRLGGKEHNLLGGRRDSRVIRLWNQQGAHRVRAAGSECAVGTAEHDRRSKALHQIQCIAAIGK

>KDN97018.1 hypothetical protein BV82_5227 [Pseudomonas donghuensis]
MRKTNDAKPDSLHVLKFLVLARELCKFSGVRSLVKATLTS

>KDN97019.1 hypothetical protein BV82_5228 [Pseudomonas donghuensis]
MPTFESMPLMNHANHSHACDGVFLAHEKFDLVSRNTW

>KDN97021.1 hypothetical protein BV82_5230A, partial [Pseudomonas donghuensis]
MLKASRSPPVPTRMKSNVALAASL

>KDN97055.1 hypothetical protein BV82_5055 [Pseudomonas donghuensis]
MCVHGVLQPFMKGALRLKTVICDTGQNIKHKNVYRVRLVYRFIRA

>KDN97076.1 hypothetical protein BV82_5076 [Pseudomonas donghuensis]
MTKRVSRFLNHSGRRNGPEPASRQRYVESGCKSMGSIEALKIVVGIVEYSYILTRETATRTVANRCCRDC
DRFALDRWQVSSHRSGACKSTVGAGLPAMGQQGCPAVNAGFRTTW

>KDN97082.1 hypothetical protein BV82_5082 [Pseudomonas donghuensis]
MGETFQSEHNARALKKASLAQHCAQGADTALIRCELRTYEHKNPRP

>KDN97105.1 hypothetical protein BV82_5105 [Pseudomonas donghuensis]
MLSWQVKRIGLVLPLCVVLRAGRCSSTARLAAAGLGR

>KDN97119.1 hypothetical protein BV82_5119 [Pseudomonas donghuensis]
MDAFTFGRVSHVGTSTVVEARKGQAVVRGRLSFTAVHYYYLQ

>KDN97132.1 hypothetical protein BV82_5021A, partial [Pseudomonas donghuensis]
MEEVIGNANAGCADVHGRRVYVRWQGVGLQRSFSGVGFSEPDAGQDVRRRRKRLGGKEHNLLGGRRDSRV
IRLWNQQGAHRVRTAGSECAVGTAEHDRRSKALHQIQCIAAIGK

>KDN97145.1 cytochrome C and Quinol oxidase polypeptide I family protein, partial [Pseudomonas donghuensis]
TPGAPCGWPMKNWR

>KDN97146.1 hypothetical protein BV82_5020A, partial [Pseudomonas donghuensis]
MPDVEGRLGTVALPVGLGCRQGDQVVRGQAGGIIRIGGVGMHYGAKLIEGDVTVGGDADGEHQLIGRGRA
ALHHASVERQVDRLISGRVGQPGGPRYHAQRIGQRASAIGAKCRAEVGGEVLRGVGRQYGFVHRQRRLRA
GGADTWPIILEHQVLPDVQGGLDAVAIPVGFGCRQGHQVVRGQTGGVIRIGRVGMHHGAKLVEGDVTVGS
DADGEHQLVGRRRAAFHHCAIERQVDRLTAGSVGQPGGP

>KDN97147.1 hypothetical protein BV82_5021B, partial [Pseudomonas donghuensis]
RGVGCQHGFIDGQRWLGAGGVDAWAVILEHQVLPDVQGGLGAVTVPVGLGCRQGHQVVRGQAGRIIRIGR
VGMHYGAKLIEGDITVGGNADGEHQLVGRRRAAFDHAAVERQVDRLTRGRVGQPGSPGHHAQRIGQRTGA
IGAKCRAEVGGEVLRGVGCQYRFIDCQYRLGAGGVDAWPVILEHQVLPDVEGGLGAVAVPVGFGCRQGDQ
VVRGQAGGVIRIGRVGMDHGAELVEGDVTVGGNADGEHQLIGRGRAAFHHAAIERQVDRLTGGGVGQPGG
PGYHAQRIGQRASAIGAKRRAEVGGEVLRGVGCQYRFIDC

>KDN97259.1 hypothetical protein BV82_5020B, partial [Pseudomonas donghuensis]
GHHAQRIGQGASTIGAKRRGEVGGEVLRGVGCQYGFIDCQYRLCAGGADARAIVLEDYLRADFGFRNRDL
EQLEATDIEIGMVSALGVDGRAVLDEHAVVVCRCGVFNLVAGFNAGDGRNPGSKADDNGRIIRRAVRVFD
HQPGADVDDANIADCNRRITHIAVDEIRARFSSGAECADFYDSAFAALQLHDVERGRPRIENVGKVDVGF
NICFVLIQVGNGHILPGTGDERKADAFLFRNHLVVGAEGGRATRLAGIDCVARRYIGAA

>KDN97269.1 hypothetical protein BV82_4818 [Pseudomonas donghuensis]
MRLDDLKCLHRVREQGGILQCCDQGLAHGRSFWVRRFESAIMAASGAGANRRRAPVDD

>KDN97296.1 hypothetical protein BV82_4845 [Pseudomonas donghuensis]
MTSNTARLEGNGRLTELEFLDYRVHGFLPFSVALGELVLANSWAAQGIGPVGSPDKTRGWEPALPSR

>KDN97317.1 hypothetical protein BV82_4868 [Pseudomonas donghuensis]
MYGQVHQHWGGSQALGLVIYTVFSNPLTMLMFSSIWLSGRGSDCAGL

>KDN97325.1 hypothetical protein BV82_4876 [Pseudomonas donghuensis]
MGWPLIVVGQHWHWIIGKTNLQCRGARLKASLRLCQFVPGKLSSDAQ

>KDN97361.1 hypothetical protein BV82_4719 [Pseudomonas donghuensis]
MAAPGITCMKIKIKIKIKISIKMKQVGPRFGISLATP

>KDN97391.1 hypothetical protein BV82_4749 [Pseudomonas donghuensis]
MSGESAGRTTLVAIFCPTLIQRRILALGHYRSRGSPRLKPTQIQ

>KDN97400.1 hypothetical protein BV82_4758 [Pseudomonas donghuensis]
MPAIEREAVAIQAPRYVQAALASSLASQLPQALRQRCGSWPCR

>KDN97432.1 hypothetical protein BV82_4795 [Pseudomonas donghuensis]
MAPSSCDFSTALCQPGRLDAAVRRPDSIEALPGRNAR

>KDN97453.1 hypothetical protein BV82_4686 [Pseudomonas donghuensis]
MWEPDDAIPERCCAAKWPPVLPGPASLRVGSCEQRMGRRTPPVLQGSAVCLRNTAKPMTEKSVFTFF

>KDN97456.1 hypothetical protein BV82_4689 [Pseudomonas donghuensis]
MTPCRQAASTDCTHITQTKDADTHRIHLMCLSRRLNKKSQTGLINQQLSGTRRRGKKMSLRRVQAYKYQV
QKSLHLNLAAKGQ

>KDN97496.1 putative membrane protein [Pseudomonas donghuensis]
MEADMFFDNVVIAGVITVGLMLMFFVGFGIFIWKDSNKRKER

>KDN97553.1 hypothetical protein BV82_4412 [Pseudomonas donghuensis]
MSKVEPKSPLTIIAIFAGIIEASALASLPFLGEDSQGIYTWFLVGFPFFLTVLFFLTLNFNYKSLYSPEI
NELTAAAAPAPAHIPLQVEPPTHEVADIDLPSTPVKEVETRHVLADMSELSTVTIAFRGPAASELIEYFV
LQALKPTRLPCNKWILSNLDTGAQITLMRQSLSKA

>KDN97674.1 hypothetical protein BV82_4534 [Pseudomonas donghuensis]
MPDARACQPAIEFHCMHAGNAKDMVYAIAFKEFDQYFAASRHDRLPVIVRETGCIGPPDSYLHNAKKLIL
AMFSNHGSSR

>KDN97681.1 hypothetical protein BV82_4541 [Pseudomonas donghuensis]
MIVVLAFFALKCERRRDKNGMKQDFSPISAIIGHCPGAS

>KDN97857.1 hypothetical protein BV82_4162 [Pseudomonas donghuensis]
MLFDGELIAVAIQQSFTYTFDLQQFIYCLKGTMFAAVSDNSFSLAPSNSGEL

>KDN97901.1 hypothetical protein BV82_4206 [Pseudomonas donghuensis]
MLLGVTIHVAGDVMSFLVALRSLPEQVSNHARLMMQLP

>KDN97907.1 hypothetical protein BV82_4212 [Pseudomonas donghuensis]
MSPLSPARPAPTASGLARPCGSWLASDSLNAVFRRVSAPLAGAT

>KDN97959.1 hypothetical protein BV82_4266 [Pseudomonas donghuensis]
MGSRSNEGFGAKLHKKQRRQLERRATLLHFSDNGPVLFCTFFQNISAFFLDVRAGILSLAEDCRRIVGHR
LKSWHARPRASARDNPQKLWITRWTTP

>KDN97966.1 putative membrane protein [Pseudomonas donghuensis]
MAIRTVLGLAAVLVLALAWWGWHEGGLALMQLGMGVC

>KDN98008.1 hypothetical protein BV82_4315 [Pseudomonas donghuensis]
MTKAEHFTLKPTAGVGQSDKDCIVIKQRPDSAICSKKPALCSERLSILRHKATQR

>KDN98038.1 hypothetical protein BV82_3920 [Pseudomonas donghuensis]
MLNGVWALVMAARLEAGTCCAPALSVRLAPGVLLLSRPG

>KDN98062.1 hypothetical protein BV82_3945 [Pseudomonas donghuensis]
MDQGVHCALQIFHGVGRPAKKHQDSSDTPLRAPTWIVVTATSLANPGDTGVFPWVCRGLG

>KDN98074.1 hypothetical protein BV82_3957 [Pseudomonas donghuensis]
MSKVRPYRSRGRRCKRDLATHMPRRPCPPDDATAIIAPSLTSS

>KDN98089.1 hypothetical protein BV82_3972 [Pseudomonas donghuensis]
MLYIMDAELIEGNYVLYQGDLSTNCVGKAQVARLPALAWPGACGLDLENK

>KDN98173.1 hypothetical protein BV82_4065 [Pseudomonas donghuensis]
MAVHQSYTQLFGLSRVNQHSFHVIPSVSGLPETAFGTHDFSRSVSGAQGVAGHPRVQQLSSPTGHPVLPK
SRRRVLLAVVSNALSQEEESAFGRGRPWGILIKTGSGRLANRPVDWTQCYTVRRLCISTLHVSLIIRVLP
RAESATGCIFASSIREPAPFMSKAGISEAVAMTA

>KDN98231.1 hypothetical protein BV82_3856 [Pseudomonas donghuensis]
MVCQVVLCSARTIRPTSRSAFCRGALLNTLSLLPPVETRWLLACPNR

>KDN98307.1 hypothetical protein BV82_3678 [Pseudomonas donghuensis]
MRNAGRLQEIAVYCIGGLYSAVDFSSRCAWLYVSDIFTLGVCYR

>KDN98317.1 hypothetical protein BV82_3688 [Pseudomonas donghuensis]
MDYRESGNSEEFEGRMFIGKVIEKNGKSLPECSGRSPFTSVLP

>KDN98327.1 hypothetical protein BV82_3698 [Pseudomonas donghuensis]
MLDNRRSWQNQPNWRRKQFESLFYCEKEKLIVFQAIMSNSQQPIN

>KDN98348.1 hypothetical protein BV82_3720 [Pseudomonas donghuensis]
MKINRIKMKTDERQFWYETCSLERVTQQTPARREINDESCA

>KDN98368.1 acetyltransferase, GNAT family domain protein [Pseudomonas donghuensis]
MTFFGQPDDLFMIANIEFTSSPTEQRGLYRYLAGYFQLPGAGVLPHARFYGFW

>KDN98370.1 hypothetical protein BV82_3744 [Pseudomonas donghuensis]
MDDQHKKIIGYRDPSQSEITGIRLQDLDRFQLMTDRN

>KDN98376.1 hypothetical protein BV82_3750 [Pseudomonas donghuensis]
MEGIVRSRLAHVVDSLKEKFGAGAMANRPRCGRWQRGIYRLKRVDRRGI

>KDN98397.1 hypothetical protein BV82_3771 [Pseudomonas donghuensis]
MALNSASVIGRVRLALSSMPFGQPLKTGGVYSLCLQAAGAGHPACAVVMQWFAEQHGR

>KDN98416.1 hypothetical protein BV82_3790 [Pseudomonas donghuensis]
MQVKAKPMAFGQCQSAKQAATQESQAEPFSWPGRLDQTVYQDEKRQDEEELIEL

>KDN98484.1 hypothetical protein BV82_3649 [Pseudomonas donghuensis]
MEHDQVFARQQQLHRYPKFEQLFKPQEPLKRRDYKGFTGWHGRCNSSLRD

>KDN98516.1 hypothetical protein BV82_3635 [Pseudomonas donghuensis]
MPTPTCQKRHSIPCWQHMREHILSLADALTAALVKQFPDKF

>KDN98521.1 hypothetical protein BV82_3584A, partial [Pseudomonas donghuensis]
MGAPQPFADKSAHKLGAEPVDAGLTRDTGVAWIATATRSIAGKTRSHKSGVHRSTVG

>KDN98552.1 hypothetical protein BV82_3559B, partial [Pseudomonas donghuensis]
AVVIQAELVSSHPIQPLSCATLPTGFPELQCTLMYPAAWCSSTKATAFGPPA

>KDN98571.1 hypothetical protein BV82_3579 [Pseudomonas donghuensis]
MLSQSGCGVRREIREKLVGLSFEYSSLDRPWRKVLLNGQD

>KDN98576.1 hypothetical protein BV82_3584B, partial [Pseudomonas donghuensis]
PTSPAYADPLWELACQRWGRYSLHIWDGLDDEREAVAIQAMLAYFRLPSPRIRS

>KDN98585.1 hypothetical protein BV82_3312 [Pseudomonas donghuensis]
MHEYRSFRHPRAGLPLGRGVMTRDWEERIQGWVSSKGVLPRRRRRCTDDEAHSRGLCFLCEIRILNK

>KDN98592.1 hypothetical protein BV82_3320 [Pseudomonas donghuensis]
MPTRGQTTGTDSAYITQSKDADIHSSLSQGSEKADRDTQLNGMVDAADAQEKAAR

>KDN98644.1 hypothetical protein BV82_3372 [Pseudomonas donghuensis]
MLVLVVQHIHQVDEALLGGDRAVFKVWQALDDPVIKMPGQFQVVDHRRGLFAEFDKIKPHRALLAQVCAQ
RALAYLQQNRRVTRLRQLQGLALFAAARPQVMTLGKTLLASSAGQATKVLLAAHFKHVHTAVDQADHRLE
TLVIERVGQQALRCVVGGHQQKYPPREQGLKQPGDEHGIANVVHMELVETQHLAVFEQFIQGAGQGIVLM
AMVKHALMQAGEEVMKVQAALVGLGQGLEETVEQPALAAPDRAIQVQAARLASFKLCRLGGHMFDDAQLA
GAQGVAAGCGLGAKMLTNGLRQGAVISRGRAQAAQGAQGRTYPGRGQGRQTGLLGAMGLKGQLERGRPI

>KDN98646.1 hypothetical protein BV82_3374 [Pseudomonas donghuensis]
MHGGRGRLKNKRRPSYRQDATLVPWQTGRCGPYSKLALY

>KDN98651.1 hypothetical protein BV82_3379 [Pseudomonas donghuensis]
MFLGVGCIALACGQSRLPMRNLDHFHNLSRTTGPLVVFLEPEIVHITLVNQPFNLVIGL

>KDN98666.1 hypothetical protein BV82_3394 [Pseudomonas donghuensis]
MEFHGGGASAGVCLLLRLPDSAPLIPRRNANARGQQHNLQYLKLARVCTDRG

>KDN98700.1 hypothetical protein BV82_3428 [Pseudomonas donghuensis]
MNPAGSHSMRSLLVGIQAVGLPPISRTTRRINIICMKNTLANGPAPSPSTACAG

>KDN98707.1 hypothetical protein BV82_3435 [Pseudomonas donghuensis]
MAGGLQLSSESDDRPFGVHPRSHLVIGSYLCLAHSSGPVV

>KDN98727.1 hypothetical protein BV82_3455 [Pseudomonas donghuensis]
MQCMDSDARHGVDSCILGVCHKSVGYKNYLPYSAVAITCS

>KDN98743.1 hypothetical protein BV82_3471 [Pseudomonas donghuensis]
MIHPIILGRGKRWFDDQAHASAFTLAQSSSTPSGVLVARYVLNGEVRTGSFA

>KDN98746.1 hypothetical protein BV82_3474 [Pseudomonas donghuensis]
MLSVPHFMFLASVLAARADQPLPGNVLTGEPGSAVFRQVKQDIARTH

>KDN98823.1 mannose-6-phosphate isomerase domain protein [Pseudomonas donghuensis]
MEHRTAAEGEAKLLMIEPRGILNTGHEGGERTASNDVWI

>KDN98831.1 hypothetical protein BV82_3559A, partial [Pseudomonas donghuensis]
MICWLHWPHRRQGRLPQWTCKHPTLWELALPAIERA

>KDN98833.1 hypothetical protein BV82_2993 [Pseudomonas donghuensis]
MHGALRKRGDECRRRLSKPSPGSHTLPWHRPGAALAARAWLPAGGTTPLT

>KDN98841.1 hypothetical protein BV82_3001 [Pseudomonas donghuensis]
MHVLVPWADGGVWPALSALSRCSTDGRIGVMCVRATNK

>KDN98880.1 hypothetical protein BV82_3040 [Pseudomonas donghuensis]
MFSSISFWSYCSTSSLYSFDSHQDAVLLGHVVVLSCRRRPRRGGVIG

>KDN98881.1 hypothetical protein BV82_3041 [Pseudomonas donghuensis]
MTMKPVALTVADHEALAELPVSDWFDVAFAPINRPSYRCQRLSKRNG

>KDN98887.1 hypothetical protein BV82_3047 [Pseudomonas donghuensis]
MGKLDNPTTMTGNYSEWTAPGTPPLSFDRHFTYSMTFLQADCPPPKQ

>KDN98890.1 hypothetical protein BV82_3050 [Pseudomonas donghuensis]
MVFSRSLLDASDVGLARPQGNQSLVAGIGDCDNSLSSRWVKFASRGSMNHAAGVGLFNGVSAQARQATAA

DT

>KDN98895.1 hypothetical protein BV82_3055 [Pseudomonas donghuensis]
MGLARPQGNQSLVAGIGDGDNSLSSRWVKFASRGSMNHAAGAGGGWHPTATTLGWGGEQD

>KDN98907.1 hypothetical protein BV82_3067 [Pseudomonas donghuensis]
MYDADHLTLVFNCRVFLRRDPELFARVARGVVAGHIDLLTAYGQDQSATLARVLSRGVRLHGG

>KDN98941.1 putative membrane protein [Pseudomonas donghuensis]
MTCYRRAKRVAFWRGSALTLLACTGFMLASSLAGAITQ

>KDN98945.1 hypothetical protein BV82_3105 [Pseudomonas donghuensis]
MPPEGQRLVGEAGQFDGLASRCEEGSFRCLQFCLALGADEDLIVDHGVLT

>KDN98952.1 hypothetical protein BV82_3112 [Pseudomonas donghuensis]
MAGEKVKQLSQMFDAPQYAQQFMELARKTGKCRDLRIRAKCEITDAQGKPVINQKTKVAKIGWTDYRLST
STHAT

>KDN98984.1 hypothetical protein BV82_3144 [Pseudomonas donghuensis]
MRDERQRCHAPGLFATGSPRPVECRKLSEVGLLACIAPAPGPVERRLPAGLQWLCRTLQCLQLRGQPRLE
PRSRLSFGQRRRTSKRLGYAVPGERSTAVTIDACPRP

>KDN99108.1 hypothetical protein BV82_3268 [Pseudomonas donghuensis]
MSTVLFSQLLQRLKGGLAGTEKGADQQRPHGSTDAGPRAWAGKAVAHQKS

>KDN99109.1 alpha/beta hydrolase family protein [Pseudomonas donghuensis]
MLAPPDQRATERLRVYLEGDGHAWATATQPSLDPSPRNLLVARLALEDPTPNAYLARPCQFVTAAGCSSA
LWTDRRFSQEVVSSLSQALDRLKQRFGNREFELVGYSGGAALAVLLAGQRDDVTQIQTIAGNLDPGLWVQ
LKGLSPLNGSLDPMAYRDRVKSLPQRHLLGGRDQVIAGEIAASFRGNLRSSDCNQRVIIPGATHESGWEA
AWRVWRSQPIDRCNQHGNRAESPRAEDRDPPRPAGSGV

>KDN99110.1 hypothetical protein BV82_3270 [Pseudomonas donghuensis]
MLKIETHRGRPVQEYDVRNRWFNLVDRSPVTDDKPFDD

>KDN99139.1 putative sulfite oxidase cytochrome subunit, soxD [Pseudomonas donghuensis]
MPFQAPQSLTNEQVYAVTGCILHMNKLTEANTTVNASTLAAVKMPNQDGFLSMRDPTAKLSPA

>KDN99176.1 hypothetical protein BV82_2956 [Pseudomonas donghuensis]
MRQFLLTGLYAQRLPGASHRTTGCFHLMTKPRGLAAWRAFS

>KDN99302.1 hypothetical protein BV82_2512 [Pseudomonas donghuensis]
MGGRFRAVIRHEALLAANQVTKRGVSFSCPLHRRQVAAGTDEAEIAAGSPCARPELINPPGMTGDE

>KDN99346.1 primosome assembly PriA domain protein [Pseudomonas donghuensis]

MAGFSLSAGTSLVSLISMSIPIISRRPNGTRMRMPGVSAWRAMLAGAL

>KDN99357.1 hypothetical protein BV82_2567 [Pseudomonas donghuensis]
MVQTCFHRHFLTRFASHQGKSFFSTTTYHPRRTNASFAGAIGEPLLHSDREAHKFTTNCRKSMSRF

>KDN99471.1 hypothetical protein BV82_2681 [Pseudomonas donghuensis]
MPRKPMQCAGREQPVRCNQGAGVAGGGAWRGVSFGYSFWGDPPKHFCFDPEFAKCLLVGCARVRW

>KDN99522.1 hypothetical protein BV82_2732 [Pseudomonas donghuensis]
MHHAGVVETMGKSMMYARRWAQSWLAIGLLWFVVAIALAPSNKVYQ

>KDN99523.1 putative membrane protein [Pseudomonas donghuensis]
MLQSVWGLALVALGMFLLLCQRRGAALVLLISVLAMPIWC

>KDN99524.1 hypothetical protein BV82_2734 [Pseudomonas donghuensis]
MLLWLAVWLYVLREIWKARDTAFGKGLMGLWLFSTLAMQFDAASLTGTPRAEWFISWLPVGLATVLVWGR
ARAPGCDKISGSI

>KDN99588.1 hypothetical protein BV82_2800 [Pseudomonas donghuensis]
MVLLEFFLVGTTTFSNCSEQAPCQWFFILEKLVNKLQRKDSRPFFIGRATVCLTASATRSSKKEQPRSAI
IATLGCGYVQPACNRMRNRTAHPWCATATAGTITG

>KDN99659.1 hypothetical protein BV82_2871 [Pseudomonas donghuensis]
MPARWRFVPRATCRGRQVSLHYKSSTAVRISDYWVSK

>KDN99726.1 putative conserved repeat domain protein, partial [Pseudomonas donghuensis]
VK

>KDN99753.1 hypothetical protein BV82_2166 [Pseudomonas donghuensis]
MCFDLLFDTPEQHNHWPGKLSTARRSIGKKGYGHLRPSALG

>KDN99807.1 hypothetical protein BV82_2220 [Pseudomonas donghuensis]
MPGVPLTHDASLVGAMKDASKNDSAQLFYAAFALPGGEGKLCETD

>KDN99871.1 hypothetical protein BV82_2284 [Pseudomonas donghuensis]
MTNEGISEGEEQSDTNTDHGYGVEQTGNDEHFNLQHWNHLWLTSSAFQELAAQQAETNGGTQSTQADQQS
NSDSGKTNYSFHLSSRLLRRIG

>KDN99957.1 hypothetical protein BV82_2370 [Pseudomonas donghuensis]
MNRLDATGHRTGRGRPEFVSVTAAGTKKGSRSHPFHLCAAVLTQR

>KDN99967.1 hypothetical protein BV82_2380 [Pseudomonas donghuensis]
MCESGQKCLNQRLIVAFSWDEGRQALLKEQLKRDPNRY

>KDO00052.1 hypothetical protein BV82_2064 [Pseudomonas donghuensis]
MAPRTNTGPGLAPDETQPELGAPAGISVDLVGEFGERHASGSSKRCSYRATGAFARNNLRAQANAATCRF
DQSISNRWQRQST

>KDO00066.1 hypothetical protein BV82_2078 [Pseudomonas donghuensis]
MPSWHGRCNTSGKGHLADSIRLMEHAHDNRRHLAAAYHNFA

>KDO00067.1 hypothetical protein BV82_2079 [Pseudomonas donghuensis]
MLLQQHPALANAAQALLRGALPPMAQNLLKALGTCSFKEALS

>KDO00071.1 hypothetical protein BV82_2083 [Pseudomonas donghuensis]
MNVRTLVQRPCHAARARNYRGLRAGVLPGQQRVSSGCGGCSTTASLASPLPQAVGAGLPREKALN

>KDO00081.1 putative membrane protein [Pseudomonas donghuensis]
MGNSTKAGKPDSSVDAWAILCLIILVVVTAVYWVSHQ

>KDO00091.1 hypothetical protein BV82_2103 [Pseudomonas donghuensis]
MGLEWPQSVGCHVHCLMDGLRQQSIKGTLARGCRYVFTRHLPHPPAQPIKPAD

>KDO00107.1 hypothetical protein BV82_2119 [Pseudomonas donghuensis]
MCSLMRLGLLCSPSPASRLPQDSVQAAPVGAGLPAMLSTA

>KDO00114.1 hypothetical protein BV82_2126 [Pseudomonas donghuensis]
MTVKVRAGARLGVQKKSAIGQGQGRNPEQVMHIPIGAGQTDGAAHRGRLAE

>KDO00116.1 hypothetical protein BV82_2128 [Pseudomonas donghuensis]
MFLDQGVVSALIAGKPAPTSSECTEHCGSWLASDGLVTFTLT

>KDO00128.1 aldehyde dehydrogenase family protein, partial [Pseudomonas donghuensis]
QGIVFD

>KDO00182.1 hypothetical protein BV82_2005C, partial [Pseudomonas donghuensis]
VTSTTTPQRRTPPSTTR

>KDO00208.1 putative flavin-containing monooxygenase [Pseudomonas donghuensis]
MTIRVAIIGAGPSGLAQLRAFQSAHAQGAEMPEVVCFEKQADWGGMWNYTWRTGLDEHGSMYRYLWSNGP
KECLEFADYSFDEHFGRPISSYPPREVLWDYIQGRVKKAGVRDYIRFNTAVKAVRFDQASRTFTVSAYDY
SIGQGIEQVFDFVGSSYSAEDIGSQCFKYGARSITSAYRTQPMGYKWPKGWEERPQLLRVEGEMAFFADG
SSKRVDAIILCTGYQHHFPFLPDELTLKTSNRLWPAGLYQGVVWEQNPQLIYLGMQDLWYSFNLFDAQAW
FARDYMLGRIKLPQPAAMAERAG

>KDO00227.1 hypothetical protein BV82_1768 [Pseudomonas donghuensis]
MPLFEPLSEVVGIVRTAHKTTFARNGVLILLPSRALQKECYFLLPIDKAISAPFR

>KDO00266.1 hypothetical protein BV82_1808 [Pseudomonas donghuensis]
MWPVKVANRCYGDARLKYSRIICDGVCVRSAHEPYPWAT

>KDO00273.1 hypothetical protein BV82_1815 [Pseudomonas donghuensis]
MSGNCPFQRWYDRRRQPRCLRTPRKGAFTTWGPMSPASLASQLPHGHCGSWLASDRRPPGTWFRVLTCVV
HRKVS

>KDO00292.1 hypothetical protein BV82_1834 [Pseudomonas donghuensis]
MLATHIAGLWTAEVIVGAGLAGDEIAVVSGNARRLHRWQSQLPHSQRYTDALWEPALPAIAPGNVR

>KDO00298.1 hypothetical protein BV82_1840 [Pseudomonas donghuensis]
MLQGGKASRIGCLSLRLAFWADDIKFLDYRRVYRRCVVLLQMNHHFMLLR

>KDO00299.1 hypothetical protein BV82_1841 [Pseudomonas donghuensis]
MIGEIGLAASGLGNPDHGWTVVQKLEIAEKTDFMAETIAANLFLTCF

>KDO00301.1 hypothetical protein BV82_1843 [Pseudomonas donghuensis]
MTLLQALEQRLRHYHVAHPAGANDQNSHALSRNDQGGCKLFANSLASPLYIQAICRPVYATRAVLWSWQA
SWCLNRLADISTQA

>KDO00321.1 hypothetical protein BV82_1863 [Pseudomonas donghuensis]
MGHARSCCAAGRRKVPAWYRTSASTADLCCGSWPCQRWAAQQPRARLMRQRSRHPPC

>KDO00468.1 hypothetical protein BV82_1562 [Pseudomonas donghuensis]
MDHTALFKSKRQINSTIRRERLKIDKAVNWDYLAGPEELLQS

>KDO00514.1 hypothetical protein BV82_1608 [Pseudomonas donghuensis]
MVGLGNYHVKTVGAQVYGSDQGQILGCGLRHGHGFSVDNPAILPRSGAGANAQRIKA

>KDO00555.1 putative ATPase [Pseudomonas donghuensis]
MDAAMLTTLAVANYRSINHLLPALARLIIRVSEQCQVWVVSHARRLVVALEQDATCNSIVLEKVLGQTRV
AGQGMLDEPAWHWPE

>KDO00565.1 hypothetical protein BV82_1659 [Pseudomonas donghuensis]
MDCGFCQIGHQYRSLAALNALALRTGVIVFLTIPTSRIENVGAGCGDKIALVRWSGSYTLIAYKN

>KDO00576.1 hypothetical protein BV82_1670 [Pseudomonas donghuensis]
MCRLRQDLCVPLDVDWRLAPKGDRLVNRNHSFHYFGLMRQRWG

>KDO00596.1 hypothetical protein BV82_1690 [Pseudomonas donghuensis]
MLQTFCSRSFQTTTAKKVAGLCQKRPKKVGPSCQNGVFPALDRRITCTETF

>KDO00619.1 hypothetical protein BV82_1713 [Pseudomonas donghuensis]
MAQWLKHNGTNRVRKADQRRSLLARYPAGLLTEAELEALLGDGSR

>KDO00656.1 hypothetical protein BV82_1446 [Pseudomonas donghuensis]
MEIPQYGDWSIEASSTDGQVVVLFARWRRIVIPSFSLPP

>KDO00663.1 hypothetical protein BV82_1453 [Pseudomonas donghuensis]
MAPLLGTLAAFLAGTLATLFALALAMHRIAFAIARIGNSAKRAEHQAQRSDGDNQQA

>KDO00689.1 hypothetical protein BV82_1480 [Pseudomonas donghuensis]
MGSINRYFGLRDVILWYSKLLTRGCRLPAITRRRFAGADARDRAFAVYRL

>KDO00761.1 hypothetical protein BV82_1214 [Pseudomonas donghuensis]
MSIRAKRLEPFQAISHLTGLNWIFGQHEESRLGSKWGG

>KDO00765.1 hypothetical protein BV82_1218 [Pseudomonas donghuensis]
MRHTKTGPADLGRRIARDYARLGGKAEAGEQPDHTEGCKRTRFDHAYSGSIRGKVTLTERISA

>KDO00789.1 hypothetical protein BV82_1242 [Pseudomonas donghuensis]
MFSDSQALKRAHIIHEKRARVGWLTKGNASKSFSIVVASKAPSLASQLPQVQCMPTLWELACQQWVFSHA
TSDTAL

>KDO00794.1 hypothetical protein BV82_1247 [Pseudomonas donghuensis]
MVRIDGDFVADCRNLDFLVSHAALRFFWCGSSSGSPERTPVERQREPRAQMPCNGAATPTSLV

>KDO00808.1 putative glutathione S-transferase domain protein [Pseudomonas donghuensis]
MYHGHFKCNLRRIADYPNLSNWLRELYQWPGIAETVNFEHIQKHYYMSHKTINPNGIVPKGPLQDFELGH
DRERLVGRGIWRKAGE

>KDO00809.1 glutathione S-transferase, N-terminal domain protein [Pseudomonas donghuensis]
MGLLIDGRWHDQWYESSKDGAFQRESAQRRNALPAAQAGRYHLYVSLACPWAHRTLIFRKLKGLESLIDV
SVVSWLMGEHGWTFDQSKGSTGDTLEHLDLLHQRYTRDDPHYSGRVTVPVLWDKHEQRIVNNESAEIIRI
FNSAFDELTGNRLDLYPEALRGQIEALNERIYPAVNNGVYRAGFATSQAAYEQAFDDVFAELDHLERLLA
TQRYLAGEYLTEAD

>KDO00818.1 hypothetical protein BV82_1272 [Pseudomonas donghuensis]
MSVVSAGAAWRSLPWGISPSVVLLTGEIMPAAAQQRLFIFSVSMLAALQSNAVTDL

>KDO00841.1 phage tail-collar fiber family protein [Pseudomonas donghuensis]
MTDQNSQFFAILTAVGEAKQANASALGVPWTFSQMGVGDANLTDPVPSPTQTKLINERRRAPLNQVKVDP
GNPNIIIAEQVIPPDVGGWWIREIGLYDAAGDLVAVANCAPSFKPLLSQGTGKTQVVRLNIIVTSAANVQ
LKIDPSVVLATREYVDTSILNVLPKNKVPGTYTKVQINERGIVTEGSNPTTLAGHGIVDAYTKTASDSKV
QAAVDGIVNGAPGLLNQLDKIARAIDNDPNFAGTMVNALAAKAARATTLAGYNIADAFTKAETNAAIKAT
VDALVNGAPGAMDQLSELAAAMGNDPNFATTMLNQLATKAALASPVFTGDPRAPTPAAGDNDTSLATTAY
VQTAINGLQAVDVSGSGTITLNATQAGSAIIYLHGALTGDRTVVLPTRWGLYVIRNATNQPFKLTVKMAT
GTGVEITQGCVSGIMTDATNTLLQQTDFNSPILTGDPRAPTPPTADNDTSIATSAFVRNVLASLGLGDVS

SSATIADMDDVSIPSGLYYVTPYTLHSPEGLQGAVIHKVYGGGGFQLLFPYSVQRVLWRRRGGSQWMDWD
ELSPKRTTLAGYGIADGLRAQSDQFPAFNSKNPGLDYKDGAIQIREAGKVAAAQSDILFAPAITFHWAER
VFKHLLMSAAGDLHWGATGKIIHTDNMGPMLFALMGAANPAAARNVFGITNTSSFGAGSGWWRDGQTGLM
IQYGTVLAQAETVYPATFQTAFPNFCLMVIPTTLNGATGDDDAFARVISWTRQNVSLKAEYGTGTGHSPA
ARNIAYLAIGY

>KDO00846.1 hypothetical protein BV82_1300 [Pseudomonas donghuensis]
MTSFDPVVQKQVAKRLAAELHFSAAELMAMSFSDMVWWLTD

>KDO00858.1 hypothetical protein BV82_1312 [Pseudomonas donghuensis]
MIAAKTGTVTFTTMAAISCTHAATVAASVSTAVAAAVATTTPFTPCVS

>KDO00864.1 hypothetical protein BV82_1318 [Pseudomonas donghuensis]
MSQDVTDVRFAELCQRFNTHLLEAGDNGWAYALEREQCGSLPR

>KDO00899.1 hypothetical protein BV82_1353 [Pseudomonas donghuensis]
MKFVTCGLSMFNVISCISNTSFHGVATSAWQQIKHSY

>KDO00908.1 hypothetical protein BV82_1362 [Pseudomonas donghuensis]
MMVLANLAARPASKDLIRSGNAVIKKLDGVAARHPTHLTAHADPATVVSASDGPTVDEIMKYRSVTR

>KDO00910.1 glutathione S-transferase, C-terminal domain protein [Pseudomonas donghuensis]
MATQTLGLLEKQLGGQQFFCGADYSMADAILTPMLDYLERVPDSAQWLKKTSKLRDYLFRMRLRPSGRNV
LTEPNFS

>KDO00918.1 hypothetical protein BV82_1372 [Pseudomonas donghuensis]
MVEALAIHEWSPPDALWKAVSVPPGQMQVTDKGVPLCSSWSASLKLRTKAFEAA

>KDO00925.1 hypothetical protein BV82_1379 [Pseudomonas donghuensis]
MQTLRLEKAKHLLKTTHPGFEVFPQPSELRKQQFVQATFLPAYWHFSGAFR

>KDO00934.1 hypothetical protein BV82_1388 [Pseudomonas donghuensis]
MNLWELDLPAIEREAVAIQAPPYVLAGLASSLASQLPQVIGYIFIFMCVLK

>KDO00952.1 hypothetical protein BV82_1406 [Pseudomonas donghuensis]
MKLGVTGHVVPQSTGRMNPGGNNARRNGARRRTPTGLRPPTAGLPNKSLRPVSGLARVMKAFTFPCRSTV
VFRRLRSLTVAGAAPD

>KDO00965.1 hypothetical protein BV82_0832 [Pseudomonas donghuensis]
MKTKFSQKLLNGLKPENKPYRVHDTNQPGLFIRVLPSGHMSYMVTWARNKNVTLGRVGKMTLEQARQEAA
R

>KDO00985.1 hypothetical protein BV82_0852 [Pseudomonas donghuensis]
MPEKWQKSGKRQASSGKNKPGGARSALSCRLKLVACR

>KDO01043.1 hypothetical protein BV82_0910 [Pseudomonas donghuensis]
MRGHGFALCLGCLSQVGPGLAVILATTMWPGRADRQRSGGIIHVMSYCKEGYFNRLRADDRRNQ

>KDO01069.1 methyltransferase domain protein [Pseudomonas donghuensis]
MAQNIYDNPDFFHGYSQPRHSRADLDTLELPKAAFDLAYSSLTLHYIDDLPRLFANVHAALRPMA

>KDO01070.1 methyltransferase domain protein [Pseudomonas donghuensis]
MIKYHRTPGTLLNQLVGAGFTLAHVQEWGPSAEQVQACPALAEEQERPMLCLVAAQR

>KDO01131.1 hypothetical protein BV82_0998 [Pseudomonas donghuensis]
MRGDEDTKPHAKDPPDHRHDGELADDLIVISSRTDCCAHSKVPRFGCFYRVKIGASMYKPCGYCNAMAPA
PILANACPTAVSSLFGGKIVHN

>KDO01158.1 hypothetical protein BV82_1025 [Pseudomonas donghuensis]
MTDYTAFSWPRIISTPLTGSNTNVCLNLPQPSKIKEARLEADRFAVLTGLGVTVKIRAVLRSP

>KDO01180.1 hypothetical protein BV82_1047 [Pseudomonas donghuensis]
MLRFGLIFGGFDVFFRCLVHFILSFVHFLLSVLGMLLSILGFIFAGFIASGQTQSSGEHNGKSKRFTHGV
FLLVE

>KDO01218.1 hypothetical protein BV82_1090 [Pseudomonas donghuensis]
MLGRRPFSCIGQRPAATPAHQCQCGQRTARNFPCTAFVVVLQPLNGGRLCALHEALPCR

>KDO01342.1 hypothetical protein BV82_0297 [Pseudomonas donghuensis]
MVHGAQLYESRRLRCTLDGRSIRALQIGAIAQRCVGGIR

>KDO01358.1 hypothetical protein BV82_0313 [Pseudomonas donghuensis]
MYAQRRALGEKRFNGPLLVDDDSRGHSYGNAPACLRIDGK

>KDO01398.1 hypothetical protein BV82_0353 [Pseudomonas donghuensis]
MTDRHGILGRKWAVLAALQSRQVYLSPGVIQWPDVGRYVGCSFFVQNFAFFRANPRKTCTFALREADIYL
FLW

>KDO01445.1 hypothetical protein BV82_0400 [Pseudomonas donghuensis]
MAFVDELRHQQMQLQEHLNLPKRRAEVAARIRDYYQSHNIAFDEALIDQGVREFFARRLMFEAPPLSGWQ
RLLSKASMARSQLYKLFLALIAAALITQCTRIAHDTGVTVEVQNSARELLRRDQDLRSNIQQQHEKLEQL
QQKAVAQPDAAVSRLLEYVRQTLPPLDQSFASDVPDFIHKTNRDSVKALVEQREAQLEQARDNLYESRYA
LATVEGIYKQRDELASLLATPTYLEGIKRFAKLKELASLADQQLLQVKDDKTLKTATRQVNELAEQILYT
SSLLEQFTLREQLLQRFKAMPLAAGDRAQLQALFAQASNSLDTQDTFQAQGQLKHLEQMLDFAAVALTIQ
VVDRTGIKSGVERCYDAAGCNRGEATDKGKSWFLVVEATDPAGISANVPVTSSETGKQRWTRLFGVRVSQ
AEYLKVKQDKLDDGHVDERMMGSKAANSLTLRFNQRTTSNPDMILDW

>KDO01457.1 hypothetical protein BV82_0412 [Pseudomonas donghuensis]

MQQVPRNARNNMPAPGEKSAKVPAMKGEYDTPAIEKY

>KDO01461.1 hypothetical protein BV82_0416 [Pseudomonas donghuensis]
MGEHRLQYTQRTGCAISRQRFACLLAIFALKAKCRSKQAVFFDP

>KDO01475.1 uxpA domain protein [Pseudomonas donghuensis]
MLNWIGVGAAAPLLTACAGLGGTSTGNDAWLDVLYFADTLDARQAGKPVVPATRLGPVTHLGQAPWMTGA
SARQARAELNPLLDANLADGVRTGGYAVLAAVLEQLRQTAGAGNCLTLENGQGWNGSGLAYLTQGGSGVE
GSQLLASDVRVSSDERVLWPQRCSGLYRQYG

>KDO01476.1 putative uxpA protein [Pseudomonas donghuensis]
MLGAGLAPEQSAALGVTPLTVIRRGGVRVAVVGITDPYAQDQKASLKQWYQSLLPAFEQARGQAELVIAL
ADVGTGPGLWLAERLSDADLLLCARGQDFWPEPVSVLQASGKRVPVLFAGSRASGAFQVRCQRAGGQWQF
SAHFHPAFDSSLTPTAQARAAQLRGLLAQQRAGHAAWLDQPLARAPQALWRRDTRGGSWDRVLHQALSAD
SDYPVLLPGLRYDSPLAAGQAITREHLISLTGSYPAPVVEAPARQVEQVLENAAEQLFGDPLLLDNSQDL
PRWLGQPWQLSYSPSGKRVSGFSEVQGQCRTFGLQYASSSGEPLWQRVEAWLSKQPADWQLSALQLPQLR
YVQGHPGWHPRELNA

>KDO01525.1 hypothetical protein BV82_0480 [Pseudomonas donghuensis]
MGAKPIAGKASSHRLVYIGPWWEPALPAIGPYLPPMIAM

>KDO01609.1 transcription factor, Xre family domain protein [Pseudomonas donghuensis]
MCGKIAHCKNLARSVVVDQRQAGNGTGILPGLPRANGRARPFVRAYPCITVFAFYLSPRHYAFYAKTQRS
IRPQSTARSPRSLPHGAAPAYRRAAIHPVAHSQREDRRSVR

>KDO01617.1 phage tail-collar fiber family protein [Pseudomonas donghuensis]
MTDQNSQFFAILTAVGEAKQANADALGVPWTFAQMGVGDANGTDPTPSRTQTRLINERRRAPLNQVKVDP
ANTNIIIAEQVIPPDVGGWWIREIGLYDADGDLVAVANCAPSFKPLLNQGTGKTQVVRMNFIVTSAANVT
LKVDPAVVLATREFVELRLSEEIGKLDSKNSVRVATTGPIDLGGLPVLDGIAIRAGDRVLIKDQVQARDN
GIYIAGAGAWVRAQDADGNAEVTPGLTVMVEQGAQLADTRWTLVTDGPIVLGSTALQFQNLAAGYSGVTA
RNTSGQIGAADAGKMFYFYGNSADQVLTLPRASELYPGAKLNLQNMGSVPVEIFPSEGEKVWVGGSNSLP
SLTMHPCTELELVRIHPTEWFAQGTGTVYRMRNLPTLPIGSNDGRLASTGFVQQELLAERGSVVPLMDGA
AVAGMSAKKSREDHRHPTDSTRAPVDSPIFTGIPKAPTAAAEATGNQIATLDFCRLLVAALVDSAPATLD
TINELAAALGNDPNFATTMLNQLATKADRATTLTGYGITDAIPIINPLPGGSLDVHGGPYAFLTSPYESA
VCQNCYWNGKDWIRHSESTPAICVVASNGAVTVRRAPAGANPIVWASNSPILDTSNILFSYLRQLPSKVS
EHGITDVYTKTETQQRITSAIAELVGSAPGALDKLDELAAALNNDPSFATTMLNQLATKANRSSTLAGYG
ITDAIPNHNPLAAGSLDVHGGPYAFLTSLSESTVAQNAYYDGASWVRHDPTKSSVALVAAGGRAMVRRWP
AGAPVGDVGLGHEVVDFGMQATEPEIDAGTTSLKWMSVAGVARFVARTVKQATEALAGLARIGTQAEVNA
GTDDTLIVTPKKLRFGVAYSLTQNGYVALPYWLGGFIFQWGYVYESQGVSDYRNFTIPFPNNVFGMNITI
EAATTGGHAGSFGHVVQVVNRNAFLWSVGGSLGGAGYGYYWAWGN

>KDO01694.1 hypothetical protein BV82_0653 [Pseudomonas donghuensis]
MAGAMSWPVTGRKRHAKEPVSSSMTTGRGEGRNHAQD

>KDO01703.1 hypothetical protein BV82_0662 [Pseudomonas donghuensis]
MTVHRPMRPAPAPLMGQRIPAYVCPGQATMAKGIGQYSEVLHATSWRLKKKGRAGERDQ

>KDO01755.1 hypothetical protein BV82_0714 [Pseudomonas donghuensis]
MKGLDAKKATKKKPLKTAKEKHAAKRAKRSGGSFLSQ

>KDO01766.1 hypothetical protein BV82_0725 [Pseudomonas donghuensis]
MHGALARAVRRTGLPSRSGGSLRQGRSARSWPHGSVQSEAGVRWAGMAAVPQGPEGENDVHQRIFPRACP
HLRERRTQQRMGQTGRRSHAWILPCRGRFIK

>KDO01767.1 hypothetical protein BV82_0726 [Pseudomonas donghuensis]
MQVSLLLHDRLFQGLRLSQICRRSARLQAARALFSQDLPEVAQRLI

>KDO01787.1 hypothetical protein BV82_0746 [Pseudomonas donghuensis]
MRNHLFNWLHDLAAALGLIPPPLQPVPIPTDEEQRKRQPRR

>KDO01790.1 hypothetical protein BV82_0749 [Pseudomonas donghuensis]
MRETEHGGDAAEARHFVAAQWLVQNLVHGRASSIRLF

>KDO01826.1 hypothetical protein BV82_0785 [Pseudomonas donghuensis]
MFIENPAEDNDNLLVVQPTVARVEAGKEQQVIFHYVGKEPLHSQRLRRVIFEGIRETAPGSTQNMRVNFG
VRQNLPVILHPKGLARNSEPWKGLVWRVDGKQLRVENNTPYVVRLAQELQLLPGQHSLDLGRNYVLAHET
ITLDLPDAAAAATTVRFQPATVYGFAVDPFEASIL

>KDO01827.1 hypothetical protein BV82_0786 [Pseudomonas donghuensis]
MIKACKLVAGPLLPAILLCTLPLPQAYADGMVPRTSVVIVEEARASGSEFVVTNTDP

>KDO01835.1 hypothetical protein BV82_0794 [Pseudomonas donghuensis]
MHEIPNLPFPSLNPEEQTPATHSAEAAQADHDGDETSSADQE

>KDO01847.1 hypothetical protein BV82_0806 [Pseudomonas donghuensis]
MHRSLLNGHRSLLKRNACLELASDGGIVRTDRRLTLPPSDNFL

>KDO01856.1 hypothetical protein BV82_0815 [Pseudomonas donghuensis]
MTKRRQRAALYRAFGLGAVCLCYRNRQLDGLQRQRQRKTAHRHPAGPLRA

>KDO01861.1 hypothetical protein BV82_0824 [Pseudomonas donghuensis]
MAIRQKNTPREGCFFCLAFTALAALQLARRVIGAPQAKKANARALALGD

>KDO01867.1 flagellar system-associated repeat family protein, partial [Pseudomonas donghuensis]
SR

>KDO01893.1 putative lipoprotein [Pseudomonas donghuensis]
MCFLIRIGGRLLLLMASRCSEQGLSAACAGAETLFQQK

>KDO02029.1 hypothetical protein BV82_0164 [Pseudomonas donghuensis]
MGNAPEDRGSRSGGPVLLIWVIEQHRGASPLLHFCRSGLAPR

>KDO02035.1 hypothetical protein BV82_0170 [Pseudomonas donghuensis]
MTYFRHVTLLSSGRSIARIVSPGSPAGIKKNLRSTWGRLLAAGPEKR

>KDO02064.1 hypothetical protein BV82_0199 [Pseudomonas donghuensis]
MVGAQGQLRAELTAQQFAAIGRVQAPPATRIEQVIGAWHPQEEMPGEVHALKGDRQATGQFQGQQGQGQW
LTAATLHDFVEQRRLGPQGVVLVLLETQGVEAAQQGLGQFAERQHRQPGADLLLQCLNLGQDLRRHHALI
VLGSNPQCRFQQGDRVTFKASQAGATWVHDNLRSSFCCSAGAAFRPVVRVLSSGVKARPILGSAPHRGAR

**Supplementary File 4**
**118 frames from complete not mapped to draft genome of P482**
>BV82_00470
MAREFDGSYQPNAKSRKQQEKDQRRMEFRRAIESYSEQRRLLQEITDYADLQTLSVWQVSAAESPRNAQQAR

>BV82_00695
METVVGLTAIAVALLIGLGALGTAIGFGLLGGKFLEGAARQPEMVPMLQVKMFIVAGLLDAVTMIGVGIALFFTFANPF
VGQIAG

>BV82_00705
METRTPNRLPFHRWAVFPVLLAQCVVFLLATLALWQWHGAVSGYSALCGGLIAWLPNVYFAWKAFRYTGARAAQAI
VKSFYAGEAGKLILTAVLFALTFAGVKPLAPLAVFGVFLLTQLVSWFAPLLMNIRLSRP

>BV82_00740
MRKLALVPIQFYRYAISPLMASHCRFYPSCSCYAIEAIETHGLLRGGWLTVRRLGRCHPWNAGGFDPVPPASSSRTSSIA
E

>BV82_00745
VVSQDFSREKRLLIPRHFKAVFDSPTGKVPGKCLLILARENGLDHPRLGLVIGKKSVKLSVQRNRLKRLMRDSFRLNQHSL
AGWDIVIVARKGLGEIENPELHQHFGKLWKRLARSRPAPVVNTDSVGVDSQDA

>BV82_04870
MNPRSACLACLERDPVALLEAALWIAAEHDESVVPAQVLAEVQALQQEVSAGLPMLPLSELAQPLLRRLNALGFQQDE
YHPLRPQSALLDKVMQRRRGQPLALALLTLELARGLSIPLEGVNFPGHFLLRVPGADHLLDPCGGRRLYPADCRELLGRQ
FGPQLALSADHLLTASPMQMLQRLSRNLRQLHASHDDYLEALKDAERVLQLGPGTAADYMARATLYQHLECPQAERF
DLEHALLLSDDPIQRLRLTERLSHLPPASPSVH

>BV82_05080
MNKPFDMEIFLAGVLTGAHATRECHIRQAKTIQAAISERWNRDNPWTWQRKHLVWFLNDHMNSNSKSTRYYYLLTT
NLITLRLGKSWNFPR

>BV82_06175
MEKSVEVKIHWVGDQFGGKRVLPHETLCYVMTELLKGASGVEANWSLVLQVEGNAEFCGNRVGFGKAWFLTKDAPF
YLLKKGYSLAVHEGSKLIATLEVI

>BV82_07010
MKSFSTLALITALAAVLQGWPALQLQAAQYQEPIDTSGVNLQPQEQNGVSYLTGGIGLDESRAIQQVQGYNLHLTFST
GPQNEYAPDVDLTIQGGQGQALLQLNQVGPMVYVKLPPGRYNLVASRNGEQQLSTVTLVPGQTGKVNLHWNQVN

>BV82_07475
MTSSPGLPHPLLRPSFPKVRARSGENPLVHIILAFWALWHCRHARPPDARLALH

>BV82_07540

MGYLLIVALIQAFSFSLIGEYLAGHVDSYFAVLARVLLAGLVFLPLTRWRQVEPRFMRSMLLIGALQYGITYVCLYLSFRVL
TVPEVLLFTILTPLHVTLIEDALNRRFNPWALVAALVAVAGAAVIRFDRISPDFFMGFLLLQLANFTYAAGQVLYRHLIAR
NPSDLPHYRRFGYFYLGALLVVLPAFLLFGNTQHLPSTDVQWLVLLFLGLCPTALGLYWWNKGACLVSGGTLAVMNNL
HVPVGLLLNLLIWNQHEPLGRLFIGGAVILASLWFSRLGTQRPLLAGQGRG

>BV82_07620
MANRIALVMGVKTRMGYISLISSNGPAIRQKSDENLRLNRIRLVTTELRTFTRAAKKIFS

>BV82_07950
MPHYRRFLGVNSVTNIEPDKGVLPVPIIRNEHWFEKFSFSRFPVIQTFVGYSMQQAETATESV

>BV82_09030
MRKPFAFAVMLAAALGLAACDKASENKAQDAQQHSENAQEKMNEAQDKVNEAAKENVEAAKDQAEAQQKAAEEA
AKEAAPAAPVTPAEPAKQ

>BV82_09440
MWKKPAFTDLRIGFEVTMYFANR

>BV82_10140
MKYEIFDVSQYQKDQVEPLGTKSKYWCSGPDGHDYLFKSIETYDSNHNLVSRGGEDWAEKIACEIAKFLSIPCADYSLAN
NGSSRGVITKNFIGKNNAYLVTGNEVLRNYSSFAMQEELAKAERQDILGVYSILRRIIRYKPVGFNSLPGVKTAADFFMG
YLMLDSLISNQDRHSENWGLIVTGKKRHLAPTFDHAASLGRNESDAEKERRLLSRDKGQQVATYVLRAKSYFYLRGVRL
RTLKAFQFFGVLTPAAALSWLEHLGKLTLEEMQAIINKVPTEVMSDVSKKFALEMLVCNKSNMMALIPFFEESLEPAFRK
WQMKNYE

>BV82_10145
MSSAFVVWQDPGTTMWEPVAKLAEIDGFFKFSYTRGALNERFVGFPRMDQLNKEYSSSELFPFFQNRLLPERRPEYYS
MLTWLDMLPGESKPIEILSASGGSRKTDNFRIVKVPERTEAGEYRLKFFISGIAYLSDDVKLEVLSLVRGAKLSCVLQPNNS
ADPNAVIVINSATNNSIGYYPHYLNEDLIRLNRACGMDSPNLVTRVSVVKVNCTAPEQYRVLCESVTPWPQGFVPFQSD
KYRLLVEEV

>BV82_10155
MRIKKAANRDLPPRMLRRVRTLKSGAQWVGYYYDGRDENGRRIEIPLGTDLDVAKMEWAKLDRKAVPKIVRLLGDVF
NRYERDVIPGKMPRTQKDNLLSLKQLRAAFSDAPIDAITPQIIAQYRDKRSGKVRANREISLLSHIYNMAREWGITEREN
PAAGVRKNKERPRDFYASPEIWNAVYSCAVSELRDAMDLAYLTGQRPADVLSMRATDCVDGHLQVAQGKTSKKLRIQ
LETNGQTNELGRLVERLLKQRKERGARNPYLIITGEGRNVSASMLRLRFDDARKFATHNARDGKDELLAEEIRKFQFRDI
RPKAASEIGDLTHASRLLGHTDKRITETVYRRIGESVKPTR

>BV82_10160
MPTNLTSEILADEEIAAITGYLTPSRQIEWLNRNGWQHVLTAASRPVVGRVYARLKLAGIKPSASSAVAETWTLDLARVS

>BV82_10165
LPPEERGTVPQQTDHKSVTALLCNAASIDTLKIKSLCCAAAGIISPVSSTTEGLIPHGRLRQAAPADTTLIASDRPHAQPAK
GYKHPGSI

>BV82_10185
MSYALRQPSFIRLKAQMSLTGRFNHALYDKVSRRTVHATVDTERSAQHLHIVIRMGSTLNSICLPLDAKSNANRVADNL
EAIANGRLDTADTGITRELLIAAA

>BV82_10190
MDIYEKRLIILKALIGENQLKDFASAHADVDASYLSQILNGHRTLGDRAATNLTKKLGIPAEILTSGSYTAADQDRITRAWL
VLGLKPPALPHPAFFNGSINETAQRVVNERQNADQLPAAPVPVVGKAILDPSGHFNALEFPSEQGEGYIDIVSPDASAYS
LKVVGNSLHPRIKNGEYVLIEPGCPYRTGDEVLVTTVDGKAMVKEYIYHREGQYRFDSISDGSPPTFLDERFVAKIHFLAAI
LKASKHLDY

>BV82_10195
MNLLDYIKGLSSAQVEALAEACKTSAGQLKQVAYGNRRASAGLAIAIDRVTAGLVTCEELRPDIDWPYLRRSAPPDLTLG
GVQAA

>BV82_10205
VADIADIGNDRAQWHLDMALAAHKAVPLKASLKDCIDCDDPIPEARRLAATGCTRCAECQELSEKMGARYAG

>BV82_10235
MTEAQQRLAEVRAAISAVLKNGQRLRRADREVQMAELNSLRLLEKQYAEQVAAEAAALQRRGRNRVSYVRV

>BV82_10360
MSKIAEERFSNRDASPVTEIFTLFNQNNIRITLELPTAVQINLNQHVGYTCAPVLHNDNAIFPIDSAEFSVTLSDVKSDGGS
RNVIANVVWWDPIRMKAKPAITFTCAPVSAGQSIQCTPSDKGYTQMRWGTTVSNGEGKQKLENSLDQLKISYSPGNP
SSETVDGPDITIG

>BV82_10370
MEWLMENLPLGFRSDHCAEARTMLGWSIEALAFRSGVSPGAIRRLEKGAGLRRVTMQALAYALEAEGLFFLPGHQPM
KGDNLRGATPCPRTRDDFHLIE

>BV82_10395
MTTIDVKSRFKAKLLRPATPDNDTSWAFVVLPKDASEQLPRRGRTTVEGTINGHHFQATLEPDGQLSHWLQVSTELLE
AAGAAVGDIVTLALTPVKKEPEPDIPADFQEALAASPEARTVWNNTTTIARVDWIHWITSAKQLKTRAKRIGNACDML
ASGKKQVCCFDPSGYYSKAFRAPEAENC

>BV82_10405
MNLHNSELTQSKQASKQASKQASKQASKQASKQASKQASKQASKQASKQL

>BV82_10415
MARIGLVLTPGFADWEYAFIAGTASPFYGIDVRFFAPAPGQFFSQGGLAVTVDSSLEQCLDWKPDVVVIVGGMVWEQ
AEAPDIGDFLRASRLAGATIAGICGATLALARADLLNTTPHTSNSADFLKQKAANYKGEALYHSCTGAMVADRIITAPGP
APVSFTCAVFEAAGLSPEVVLQFRSMLAAEHE

>BV82_10420

MAHLDHLANYVSAILGLSSPRIAHLEHSNQETWLDQGFTVSAEEHIYAFDDGALIKRLAEQDNFPGDAACAECWISYEI
VRQPLDKTVSPCHISFNSACREAYWLRYFYA

>BV82_10425
MNVVSLVIAAVLSFSSVAAFAHSGGTDAKGCHTNHKTGGYHCH

>BV82_10430
MKKTLVAVFACAFVMGTAALLPADLSPIGSAYARGEGGGGGHGGGHGGGNGGGHGGGMGAGNGSHGGKSNGAS
FGGDHAGKATRDHGVSGNRYGSDRNDDNGHGTSTSGIAHSKDTRGLSKSTAISATTPGDHNAKGLSNAVRSSSKNDR

>BV82_10435
MKAYKTCSALIATAFLFLHGCENPEGHIYQANRCAVAYSMGSNVDPSVVTNAALETGQYMRAHGINKSAAELTAMTD
KVKDEIMGTPDAPYKGWEGRADRISESDFCKKYLSSLQAQ

>BV82_10440
MESDENAPGNRSKQNVTRTTTNETGHDSRRDEPEVPLPPDDETPVEQELSDMHTNHSQPTERPQAVNKGRTDQPDA
DSQMNDQDENKEPRGVPASDPESGA

>BV82_10475
MAFSPRERTALLALKGVGPTVITRLEQMGIESLAELGKADVSDILAHASAALGSTCWKNSPQARAAITAAVDLARSTPEH
SLAG

>BV82_10695
MECFSEIHERLNNRRFTVANNTQGLSGNGTVFHYLVDGGTITGTYHGGRIRMGSQVGRVTGSNTIELLYHCITTADEILA
GWSRGEIGQDHHGRTTLHFVWGWLSGATGEGESDYVERVDSPVDC

>BV82_10715
IAPVGHAGQTGTAYWRTCQFGSARVRMVEYSPGYLTDHWCWRGHILLCLKGRLNPPPKAVGRQRLLPAKSDPWAN
DESSICCIN

>BV82_10720
MPFARISLHRGKSAEYLQALSQGLHDALVESFEVPKTDRFQAIHQHEVGELIFDPHYLGGPRSHDFVLIAITAGRPRSVET
KQRFYRGLVEKLGRTTGLNAEDVMVVITTTASDEWSFGGGRGN

>BV82_10745
MENVVELHEHLDQVRDAETFLRFARALAADKSTSIDWENTTIEAFLEGAISWAESSGFGLSQGLLPSNSWRQFAVFLYC
GKIYE

>BV82_10750
MKVQKPKGKPASDNLQPSARSEEYWELIKRILLAGVMLCFLILVIPSQWRVVYSATSVFLIAIGVGIRMAQINHPGKYEFT
EFRSDAVRWYENLNRRHQLYLNVTMCIPFLGYLYVLDANGLFVPVAALFLIYCLGVVSYDVCRIYAIICETFIGKGVISVIFV
IGSNLALSISGKIIGGMTHVPPTTFPHTLSFLAILAVPFLFVAAGTVFIFIASTMAPLIIYGSSFAKKAPRFVKWFFGVELDGS
RSRYIVATLIFQVFFYVGIGTLTPAVMLAVINKYSQQIELTIGRSIYEFDMYPGTECKAAFGHRQASLGDENYILASEQSAG
VVFEPPRKCSL

>BV82_10755
LTTHALNYEVMIADLKQVADKFISVNHFKVKNAEVILQISDSQEINARVYLSASDQYIVRLYRGVLESCLSFVEDTVDSYYP
LVAAQTHLDRETFLRATCSCVTEYMFLHEYCHVIRGHINYRDFRKPLWVERPELELQDIRFLECDADIYAANFLFARTYSV
FKDPKVAIRLCDLMQCYVVGIRSMFEMLYRVSEVEDLLHEESEHPHSLVRAYIAVCMGVSGPVSECLDEDAEECRALAM
RELLSYEAFLYKDRPIDPGFLQSNFERELNIWTTREQDLVFFNLLKVQKVSIFTKAREWLRRSKRFTRDSKIQTQP

>BV82_10760
MAKRYELSDEAWAVVADLFTETHGRGRPRLSDRLMLDGVLWVLCSGAAWRDMPERFGPWSTVYQRFRGWRNQG
AFDQMLKRLHLRLNEQGLIDVQTWMIDSTAVRATRASSGAGKKGGPDEPADHALGRSRGGLTTKIHMLCDANGTPLR
FLLSGGQASDISYAQPLLDEVSIPSSQRGRPRKRCKWLLADKGYDAEALRRYCDQYRMQPVIPLRSMKRKPKPGLPRLF
DRPKYRQRNIIERMFGWLKENRRIVTRFDKLAKSYAAMVSLACSMRCLRHLFSYRT

>BV82_10770
LRNYEDMIVIFLDILGSREMSSFEEKYKIHRIFHESVKQSQDLQGTKEKEHVAYTRKLFSFSDCAYVFYKYKPDVKDSKKDL
SKLFQVALLNTSVMMLRLLNEGYLVRGGVTYGSAYFDELSFFGPAVEDAYLIESTKAVTPRILIDQRFGEKIFLHEKRIYGE
VFGPSSPHYKFLPKRSYIPTIVSPDQSEYILNVLYILEMEGSIQLGDALITHCELKDNVTSNLLIKIEKHKKDPHITRKLLWKK
NYIESSILSLESEGKSFTRLV

>BV82_10775
MMKPMNLHTDYGLITVTNEPTHSFNSTDNRRCYASEIPLTDSIGSNHGVSLNGELIVVVGAGGGATDVHAHSALVIEDT
MLLAVGNHIACISIVHPYRLLWSLQVDVATCFGIYWQSQRGALIAHGELAISRFSTDGRLIWQAWGADIFTEGFDLLHD
HIRAVDFDQSVYRFDYDTGELLT

>BV82_10780
MTKPVLHLMSGKIASGKSTLAKSLAAEQSAILLSEDHWLSRLYPDQIKSVTDYVRLARQMREVLGPLVIDVLKAGVTVVL
DFPANTPQDRQWLRDLADAAEVSHCVHYIEVDDDTCRDRLHLRNNRAEHEFAATDAEFDLITSYFRAPDEGEGLQIEV
HRL

>BV82_11350
LDWVKSVIIKMTEVMMSDCSDLNAILDRINKRMAPVSKDLRHADNRCVVDCTGELLLNLVEQKYGRLTLEDHEICTRPA
FLDSPRLRVGVTVSTTDNPDVVNLACIGTLVETWEQFFYELAHESVHLLGPADTNVVPVAAIEEGVAVKFAEDFYAEFIF
PVTGVKPLHSPLTSPRSVYFKAHQRVSKMPDRLLKDVRREFENFHLVSNPGFYDMAKNLLHWRRRRC

>BV82_12290
MNKNQVKGRVKKIEGTIKEVAGKAFGNEHLEERGKIEKTVGAIQAGLGDLKEDLNR

>BV82_12500
MAFHPQAEKGWGSNLDVGVEQEPGQAASPGLYGVLILEGEKYPRRLGEIDRYRVIFTLVSPASARNHARAQPLHWRIQ
QRLAAYAVARRAP

>BV82_12950

MAFHRLTRTHPRLSFSAVVGSLSGWLIPADGLVQHILTGWNIGVWLYLLLVLWLTLRASPEKVREVACIEDENAGLVLFT
VSIAAIASLAAVTLQLAASRGLVGGSQMLHYVYTGFTVAGSWLMIGMIFSLHYARLFYTAGSKEPLLRFADGELNPDYW
DFHYFSFTLNVAVQTSDVGVADRQMRRIVLAHSLVGFVFNTAILGFTINIAAGLLN

>BV82_13120
MSKTAKNTFTVAGGPMTALGIAFAAVGASGQPAFGYTAVGLLIPGVALIATGIWSKRRQA

>BV82_13130
MSEEYVLIQLAEAACGANGMLWQQAYHCSVRVTQRTSKQWINDPSSFGQNEQHHFLPIAAPLYMAWMIWKGPRP

>BV82_13220
MSPHVLIGEAISQLEQCYTLRADKRLEALIVNHFTAGTLDSDEFKHYCERVRRVAERRKEVA

>BV82_13240
MQVSKVIGGAVLAGLVALSGCASIVSDSKPKVGVYAVPRVTSFTVTNSAGRVVHKGQTPNQVILETGRGYFKRETYTVT
FHRDGFPDDVHVLRPTVSGWYWGNILIGGLIGMLIVDPLTGAMYTLPDDVTSNPIPLAPAQASVN

>BV82_13250
MTMNEIKAGYVIKHTKAINYISEQVDQANIGVMQSSTGATKIAMIFSRDIIEISHETLVEVPSVPGGLQPQVQHDALSTS
RVQYANLTMTIEAAESLILGLRQTLDGLKAQSSES

>BV82_13265
MKKIPLSKYLEEHGTQAVLAAALGVNQSAISQMVRSGRSIEITIFADGHLEASEIRPIPARPKRSAA

>BV82_13310
MEIDENAPGNRSQQVVTRATDNETGHDPRREESEVPLPPDDEAPVEEELSDVEANNSVSSEHPESGSGGTPGVDPQM
NDQGDDAKEPRGVPASDPESGA

>BV82_13315
MKVNVKCKSCGSDKFQIPPRPNNNSKVTCGKCGAVETYGNIMKAVGDKVTEDLKREFGKLFK

>BV82_13325
MTNVTRIHHALPLSNAIKEVVVDLDSTIAKAIDAAKAAGLPQGLIVSLLHGHAYAQTSIMVI

>BV82_13330
MADKYVGKIMGVGTDNVEYVINIYQDETVERSSHGMVRHEGLKHFEMQFGGDVKKISETEFEIVATGVRVTVHDTQS

>BV82_13360
MSHQENADFLRRYIKGDVKALQGRDIILQGDALNPHVDDFEAVIDIGKYQYVGYTYWSEEGNNTAMYQALVMFAYGR
ESLLAEQEKSS

>BV82_13415
VSNNAICCPQCQGQQVQLLSVIHAAGTSRIHTTHQAQSGIGPRVTVETTGRHQTDLAASVGPPPGKRLLGPVILTGVGA
IILYDGLKLMNSYWGVDWSRFFIGVTFIAIGVVGFVRHWKFNVAQYDKLEEWRRTWMCHSCGTRFQPS

>BV82_13425
MAIETFSWATERGEKPEIKYRVRESRFGGGYKQTVGDGPNNKEDSYPVTITARKQKALKVMEFFDRHAGAKAFLWTTP
MGQLGLFTCNDPAPVPIGGGLFKITATFERSFHP

>BV82_13445
MAQVAIDTKKYLSLTDKPSEDHWQKVYGIGSKVPASGIYRCTTCGDEITSNKGDQFPPQNKTQHPCQDEKIYWELIVM
TQTKGSGR

>BV82_13475
MRIWGADGALQIDENSFTIRVALSTLVTFEAGAAKGSQDFAVPGVGPGNGTAIVIPIGPYSASQTQFETEMLDGVARVY
NHTRTYASSFRSSGTMRLIVMRWS

>BV82_13565
VIKKIILLAMMATTIASCASVGEDKYSEKYIRSHVIENKTTKSEIQSIYGTPDEQFAGSNTGSSWVYRKRDGYKSLDSLSSFI
PGAGSVLSSAGIAQNNAEAITSISNKASGNAELSGNVLSFSFDAKNVVISWELH

>BV82_13595
MNRLLMLALLATLLSGCASLSASSAWKDRPVGELIDFFGVPRQIMLAPEEDMVVLRYVRDSSYVSREAAGTYTGPQNG
QLVHAEYWEDVQHSGSCEINVFVNRARLVQKVRTKGRCVAVEMAPAG

>BV82_13600
MLKKLTILGFTLALTGCGLLPPPAPPAPPTVEHSIASKTEIADAKRKLLRHISDPESAKFETIYKFKGRFASGKSYEGVCGYV
NYKGAEGGYEGFTPFMVVGEVVSYYGDHLPHNYHILKSFCTKSRSS

>BV82_13605
MHHTAIGPFDITLNPEALSSAAEETGLGRLSLDKQFQGDLEATSQGEMLSFRSSTQGSAGYVAMETVHGTLHGRSGSF
VLQHSSTMNRGTPVQSITVVPDSGTDALSGLIGSMVITITDGQHSYKFDYALPDHQI

>BV82_13615
MLGLMPELIEQIGPSVHTSWQNNGNGIDPLVLIWPLLHLKAGLPEEV

>BV82_13630
MIDKVIPTALACFCLLGTSTAAAWTKVNGRIQLQTEEISHCETLALDAAAVMVSRQWNLPVDYLDEDEAPNQAREEAR
LRLLAAAKSYAVRESENEKTEELKAFVNVSRRKCILDYIDSTPPTSIK

>BV82_13635
MPQPWYSTRPVPVFMTVFYCFIVLIQAGLWMVNEGFEQTRQGDISILVLLNVCTVALLMCVGGLMLLGRKKSSSLCFIL
ALALGVTFSLSQMRSGSIVHLLTLPMFKEYALVFGIWVYSLQLRSGSYFYPARSAS

>BV82_13640
MNQKLTCKKCGKETEVQLRHDSKTDWQVFNCQLCGALHVEESYFKAPGAPAQFRFRLADES

>BV82_13685

MSNFEKGDLVVLNSGGPVMTVRYVDNDISEGYSVQVTCDWFDKAEQHHTATFNAEMIAKGDGRLKGSTL

>BV82_13730
VPTAVIVKRTVLFGDCDPEGIVYTPRFSYFVIEAVHEALAVWLSGPGLRTLLGFDLLPPARAFSLEFLHPVTWDDELSMKV
SVARVGRHSFTFLVEGRLASEVVAFTASLTQVCISPVTRAVIEVPEPLRALLQA

>BV82_14250
MTEPIHDYDPADALVDPEAVDVFLADAYETGDAAHIVEAIAVVARANIRIDAADEPCKP

>BV82_14275
MNFFDTPGTRADFEERMGVLAELCRTGRMRFFEGVGGMESLTRVRYLPNGRVDLLSIDESARLQANSAHQMPNFAA
LLAADEAQSPTAVDS

>BV82_14300
MTTLVTAREYESYDEKIRAQAEMLTSLLSAGLQTQDAADKAVLALSRLQQEVHRQISSARNDLSTLADATATEAAKLLTE
KFDQANEAAAVATKRYEKAGRLLGVKTFVTLVGALAVIGAAAWFLASPLLPSHEEIEFRRRQIAEMEAKAAVLAKKGVN
LDWDQCKTGLLNRTKLCFRSDGEIYTRKGTDQHYAVPHH

>BV82_14315
MPNKTPENAFLARLRGRKSTRSNLWLVYSHKTRSDLILESHQELAHWLLHLEFDPNIVNFWIPGADELYADQRGSRKTR
PDVIATSASGGMEWHEVKAGYVEENTVSEQIIVQRQMATDNGATYRLFDESTRLTRRYELMPRLRLMHFVSVSRSQSI
DHVRQALLNYVSEAQSGTLGEMLQAYSALPPTHLIASLIQLAIEGRVTLDIEQTTPTRQTRWSARASL

>BV82_14395
MFSHVTIGTNDLARATIFYGRLLAPLGITLLALKHNPERALFTQSTTDSGTAFCIYSPLNGERATPGNGLMVAFEAQTRA
QVDDFYAIAMANGASDEGTPGLRPQYSPDYYGAYIRDPDGNKICCVCHSAD

>BV82_14420
LATWVASLPRHWPESLTSQQCDVGYGLVCDAEGKLIDGLQLRSANQEHGVASFEEGQKVEIEQQFPIGTRLRILPNHAC
ATGAQYQAYEALEENGEISHWSRLNGW

>BV82_14425
VIKNGISPGFSWGGEIFQTPVLTFQDGSQGQGRWYSSGGQLSAFYDDIQQEVHALLVGYLTYRATEEPEVAAVEILAQ
MSRHERVACERGKMQLEWYGKVLEEAGTANQQSQVGA

>BV82_15425
MTKPVLHLMSGKIASGKSTLAKSLAAERSAILLSEDHWLSRLYPDQIKSVTDYVRLARQMREVLGPLVIDVLKAGVTVVL
DFPANTPQDRQWLRDLADAAEVSHCVHYIEVDDDTCRDRLHLRNNRAEHEFAATDAEFDLITSYFRAPDEGEGLQIEV
HRL

>BV82_15435
MEKLSNDFSGALRTFAYFMASGTHYMLKNVEYLDIYGSEPSAIEMVFAIFANVIEMDNEGNVLNFTHAQRRATDYLRSY
CDPSFKVTPPFEDWETELYGPPSLGR

>BV82_15910
LPRDVYAFTGGSLLGESPTVGGTVGLTGTDGNVHDLLRIPTLALANC

>BV82_16075
MNAQTCACPGCNCKLGEHPVMSEGRAYCCQACAEHHPSGKPCNSSGCGCAASPKR

>BV82_16090
MANKGTSNPGNFANDKQKASDAGKKGGQASGAAGQPRKPDMDKDQPRKPSGGGRS

>BV82_16470
MDDPFLPQRLFALAALVVAAIGGCVFVFILLSPEGPFPGLGSALALFTLGVTNGLCLPLVGTYWWLAGKPAGAATLLILQ
VIGLIAFIGWFAAA

>BV82_16500
MKLGIAGLLLVTLCGCATVRTVTDESRAVDDLAKWETNCHSIPRAYSGVAYQFCNLNGPARSTAHWAPAPILLDMAAS
GIADTVMLPYTGYRQFKHGNTPVRRLEY

>BV82_16505
VRARYLALPPLMAFTLYTGWTLLITEQSLLAFGLELMSRPDTAQVVVDLYLMAALACMWMVTDTRARGGALIGVLPY
MLITLVFVSIGPLLYLVIRGAERKKG

>BV82_16540
MLVVLMVVISRSYGYRGADIEVMRGVAVHQTTGNHPKITLRDLDSGRLKQLDSKLLRPSDLFRKAQGKEIELSFVGSSIL
HCKVEGVDLCSPMCLSGNECEEWASAKETRKARLFVFMWLTVAVVSHVMLMMNSRRHPRSL

>BV82_16555
MNLHNSELTPSKQASKQASKQASKQASKQASKQASKQASNQAIIS

>BV82_16690
MRQITVGTKPVRGSIRRLRALERWALGFHGSCYPQSEHKEAYKNWKIPVHSGLVQGPRARVENQAFCIQQLLEAANHL
SKAADRSVGYYRVACLLVWPWVHQSEVTVFYDRDYYLGFLGQANTLESDLLSQTLALHLPGNFIEHGHDVTQPEDEVE
VQWWCIGEPA

>BV82_16695
VSLDLYLEADKALTMFTLRQAMEHAGAFQINMAECGLDAAFSSGVRLTGDGAVTDPAIYAEDTKGIDFQVAMRCSIRI
KGPEPDGQSAMQDLNKIAESIAQTCASLFLITFQFEETLYWRDATGLHSALREDATRHD

>BV82_17320
VARSISGRLGLALVLVLAASTALAANTPCSGRKGGIAGCDGDLFLCNDGSISGSKKSCTAYMGGSAARPSLQPLRGASEG
CGCGSGLFCTGPRGGVYCLTPSGKKSYRRR

>BV82_17405

MLMKNLKSLHESMKSQTNAHGDVIELQRFRSVQGAAVFECIFSTGESPYKLSLTSRGTESHPKSEFFLFDVSDEYTIPNYF
HGDTHPRLLELLKTLGGLTGNRLYPADFLAQLDANVPVLATVPAIPSSDYMLTNRPDLTDERDRPYWSHWSKPRSKAD
GSPGQVSSENQEKTAALLGSAALTYSRTMRMSSCWSPVPTNKNWRQPS

>BV82_17420
MFGFIANKAKALTEVVTEKVSSASDAVASAKGAVLESVGEGATAAVSRVEKHWPAVEKVLVDGLLTVTHDRLKDDEFFL
LAVEKAFELLPAPVRLVLPRSSIIKHSLTHRDSIVEKIAAQRAVRSLSSEPKLIETDV

>BV82_17470
VTGIVKKQLGAASRKLQEKAGGELKPRMLAQSFRAIAGRLTGWAEIFGKYALAVNLRGFFPLRV

>BV82_17640
MLKMLLVYGHLIATCIALGRVLQADQKLWRWRKDVLSQVQREYLEETQKIVTLALLALWGSGMLLVLQGYLYEGSAYLL
NQKLWAKVSVVALLSLNGALLHRMGFPLLQKTPFVLLHNSARTRLALLGALSMSGWLFAAFLGVARAWNHVLPYLHV
MGAFTLFSLAACAVALIAANAAGSMMTRLTRP

>BV82_17860
MQIQRTYEYAGHFGIVFHVSQLVVLPDHRARLRELHISEAKGAHYEVNTMLVALYDRMQAGNRACEMKVAISQPLNR
RQADKLNDQRFYASGVVGSAAGGPVGLLTKAIVGTAVTAITTYLAKRKLPTHHAGDVIVSIQASVNGGIGPQQSSVSTII
KNQ

>BV82_17870
MSEVTQLCYALILIAFLDHCLKPYLIRKWLGLFLCAAGITLLFTPGGTYWEAMDLPLAPLLVFSAGMTLIIGRRKFQPDE

>BV82_18045
MRRLRKSLGAAFAVFMGVSGGGLPGLLSAKAEANVANITFEYTGPRPGYRPDEFVNTTPPAEFCSGLVHCTRYNEHKYT
FNFPASYNKTTFQQDPDLRNRYFFKLPDVTTLHLVNEHGDSFRAQFKPTHVGLRIRSTRNLEVDYPNNGGCRGVASRN
PASLNTEYLGAIIDPSLGCWIGERHHSPGARGEHWTHVMGISFDLQLPSATSLKNGVYRAQLPWRVAFPGGGSLTPVT
VPTR

>BV82_18210
MSILDGVSLLLAVALMIYLLVALLRADRG

>BV82_18320
MDANAFAKALKTSISEVKSQGHETVNCDKLITYIDRVVEATSTDNNPEENSEVLKARLQLSLDAHKHNLSANLEMFKSII
QSGQNALKTMLLLNGGAAIALLAFIGKLSDNNSSAIPDFAYSLTAFVIGALAIGITSCLTYLSQMAFDSAVKWHNNVATT
LHATCGALGLVSISAFAYGTYKTYQAFLCLATCA

>BV82_18715
MRRLKRDPLERAFSRGYQYGVTGKSRELCPFTLPSVRQAWINGWREGRGDNWDGMTGTAGIHRLNELHAVG

>BV82_19735
MRNTVLLYLLLSLFASASLPVNAATQPMAESPPTAVIERQAGLVNLNQADVMTLQSKLTGIGRSKAEAIVAYRSEHGAF
ETVDELLEIKGIGKALLDRNRDKLTVK

>BV82_19765
VRLGELLGRNANNLDVFRMVAACMVIYGHAYAIAPQAGYSDWVERLLGYDYSGLLAVKIFFFISGLVVTNSLLSRRDAIA
FVCSRFFRIWPAFMLVVVVSALVIGPIYSSLSLVEYFSSRQFKVIDYVTSNFLMDIRYYLPGVFFELPYEAAVNGSLWTLYY
EVGAYLGLLLMFLLGIFRDKFLATLALVFLLVSQHFVNADAMHPIFHQPILVQCFAFGALLALYKDSFELHSGVVFFCWIV
FYACAASAFSKYLLYIAIFLTILYLSGTKLLLAIKPKTDVSYGIYLWGFPVQQMLAYHFLDWGVRFNQVAAIVVCIGLGYLS
WYLVEERSIAFGGRVAKNLKARVDRLVSRLSLKWKSQSA

>BV82_19800
VRKLSRLMIAVLLAALVLITVTFILENQQEVTLSFASYSLPSLPVSALLVVALVAGLSVGPLLVMMTAWRTRGPRKKPLSG
A

>BV82_22265
MDFEKLAVWQRSKDLTVAIYRQQRVCRDLGFKDQITRAALSIPSNIAEGMERRTAKEKVRYLCIAKASCGELRTQIIGGE
VGYLPKPLSNEWITETRELSRMLSGLINKISD

>BV82_22750
MSTQECALIHLTLFKHSAKSFLAALDEAGIPHGPVHMFSSAPQNSGLVEAIAALSDAMPWNAIAKIVVAWIEARKSREVI
IYTEAGESFHAKGYSASEVQKLLRISTNVTVIDTKPVDEQ

>BV82_22805
MTSRLNPEDQRRVDEYLRAPQHQVERKPFRPWLLLLLVVVVTVALGLLSRLLSHLVL

>BV82_23185
MLETVIVVLHLLGALGLVVLVLLQQGKGAEAGASFGAGASNTVFGSQGSATFLSKFTAILAASFFLTALGLGYFAKEKAQ
ELTQVGLPDPAVLEVKQQKPATDDVPVLQEQKSEATKTGDVPPAQEQK

>BV82_23525
MGESQQQATKPELYERLIDRLGLALEVARTSVRLRDEVPAELELRGLSRAEFEVIKAYLESLPEGGHGRVGALPQPETPTS
ARIVWLKDKRSATTAKSRTLQFK

>BV82_24090
MKVRASVKKLCRNCKIIRREGVVRVICSAEPRHKQRQG

>BV82_24105
MATVKVTLIKSTAGRLPNHKLCVKGLGLRRIGHTVEVLDTPENRGMINKAYYMLRVEG

>BV82_24645
MWTKPAYTDLRIGFEVTMYFASR

>BV82_25690
LACGNGSHRDPHPCLT

>BV82_26240

MKYIPFLVFLIFSAPAAGSLYIEKVAGTQYYGMFGDMDNIRAKEIYKSHVPSTMTAVWPSKNGIWIIFVFPGVAWQDPI
YLGQNLLPTDNADAVLGQLVNSQLAGGIIMPLVTTASFKSLIFMVCDYKTNTMAMTCKMATTAGGEITPPPPEEETLSC
SITGTIELHHGSLTLETVANNSAQTAALVSCNRNASVSLSIEGMVKLNGVEGLYSQLTVGGAPVGQPYKFTAGTTYTPVD
FKSVLRTVGTVTPGDFSGSAKVELAFP
